# Catchet-MS identifies IKZF1-targeting Thalidomide analogues as novel HIV-1 latency reversal agents

**DOI:** 10.1101/2021.03.19.436149

**Authors:** Enrico Ne, Raquel Crespo, Ray Izquierdo-Lara, Shringar Rao, Selin Koçer, Alicja Górska, Thomas van Staveren, Tsung Wai Kan, Dick Dekkers, Casper Rokx, Panagiotis Moulos, Pantelis Hatzis, Robert-Jan Palstra, Jeroen Demmers, Tokameh Mahmoudi

## Abstract

A major pharmacological strategy toward HIV cure aims to reverse latency in infected cells as a first step leading to their elimination. While the unbiased identification of molecular targets physically associated with the latent HIV-1 provirus would be highly valuable to unravel the molecular determinants of HIV-1 transcriptional repression and latency reversal, due to technical limitations, this has not been possible. Here we use a dCas9 targeted chromatin and histone enrichment strategy coupled to mass spectrometry (Catchet-MS) to describe the protein composition of the latent and activated HIV-1 5’LTR. Catchet-MS identified known and novel latent 5’LTR-associated host factors. Among these, IKZF1 is a novel HIV-1 transcriptional repressor, required for Polycomb Repressive Complex 2 recruitment to the LTR. We find the clinically advanced thalidomide analogue iberdomide, and the FDA approved analogues lenalidomide and pomalidomide, to be novel LRAs that, by targeting IKZF1 for degradation, reverse HIV-1 latency in CD4+T-cells isolated from virally suppressed people living with HIV-1.

**One Sentence Summary:** dCas9 targeted chromatin and histone enrichment for mass spectrometry (Catchet-MS) led to the identification of IKZF1-targeting thalidomide analogues as novel HIV-1 latency reversal agents

## Introduction

Combination antiretroviral therapy (cART) effectively blocks HIV replication and has significantly reduced AIDS-associated mortality. However, cART is not curative, has side-effects, and, due to high costs, its global roll-out remains an ongoing challenge (UNAIDS fact sheet 2019). HIV persists because, subsequent to stable integration into the CD4+ T cell host genome, the provirus can remain in a nonproductive latent state, defined by the absence of HIV-1 gene expression. Because of this reservoir of latently HIV-1 infected cells, interruption of cART leads to rapid rebound of unrestricted viral replication, necessitating life-long treatment (*1*).

Strategies for HIV cure aim to eliminate, inactivate, or reduce the pool of latently infected cells such that the patient’s immune system can control viral replication upon cessation of cART. As quiescent memory CD4+ T cells, which constitute the main cellular reservoir of latent HIV infected cells, have a long half-life (*1*), pharmacological approaches aim to speed up the decay rate of this infected reservoir. One such strategy is to induce viral expression in latently infected cells using latency reversal agents (LRAs) to increase viral protein expression, rendering the infected cells recognizable to the immune system or susceptible to viral cytopathic effects for elimination (*2*).

At the molecular level, the expression of the HIV-1 genome is determined by the activity of the HIV-1 promoter, which is controlled by the 5’LTR chromatin landscape (*3*), the engagement of sequence-specific host transcription factors (TFs) and associated cofactors (*4, 5*), the recruitment of RNA polymerase II (Pol II) and its efficient transcriptional elongation (*6, 7*). Co-transcriptionally recruited host factors subsequently mediate post-transcriptional processing of the nascent HIV-1 viral RNA template for its efficient splicing, export, and translation (*8–11*). Host factors mediating these regulatory steps are thus the main targets of ongoing pharmacological strategies to reverse latency (*12*).

Several studies have shown that pharmacological reactivation of HIV-1 transcription is possible in vivo (*13–16*). However, due to the complex and heterogeneous nature of latency, the currently available LRAs are incapable of reactivating a significant portion of cells carrying a latent provirus (*17*) and have limited capacity to induce viral protein expression (*18, 19*); thus they have failed to significantly impact the reservoir in patients (*13–16, 20*). Additionally, LRAs target host molecular complexes with widespread regulatory functions are prone to non-specific pleiotropic effects. Besides, as viral clearance is a necessary step towards HIV-1 elimination (*20, 21*), maintaining an intact cytotoxic compartment is of utmost importance in HIV-1 cure strategies. While being tolerable in vivo, LRAs can cause significant cytotoxicity to CD4+ and CD8+ T cells, and global immune activation and may lead to impairment of T cell functionality (*22*). Therefore, there is an unmet need for novel LRAs that are able to significantly activate HIV-1 transcription in vivo while maintaining an intact immune compartment.

The identification of the full repertoire of regulatory factors and cofactors involved in silencing the latent HIV 5’LTR is critical for a more complete understanding of the molecular mechanisms mediating latency and for the development of more specific and effective combinations of LRAs to deplete the HIV reservoir. However, due to technical limitations, unbiased proteomic analysis of host proteins associated with the latent or active HIV-1 promoter, in infected cells, has never been conducted.

Locus-specific strategies to identify in vivo chromatin-bound protein complexes at a single genomic location, such as the HIV-1 promoter, are largely based on a reverse chromatin immunoprecipitation (reverse-ChIP) protocol which relies on the targeting of an affinity bait to the chromatin region of interest, followed by purification of the bait and proteomic analysis of the proteins that co-purify with it (*23, 24*). Such techniques represent an enormous challenge in protein biochemistry and proteomics; in each cell, the number of protein molecules bound at a specific regulatory region is exceedingly lower than that bound to the rest of the genome (*25, 26*). Consequently these approaches require high amounts of input material, which despite rounds of purification lead to high signal to noise ratio and low relative abundance of the single locus of interest and its bound protein. Another caveat in locus-specific techniques is that a large fraction of the exogenously expressed bait is present diffusely in the cells and not exclusively bound to the locus of interest thus contributing to the background signal and affecting sensitivity. We, therefore, reasoned that a biochemical purification pipeline specifically designed to eliminate non-targeted bait complexes from the sample, would constitute an effective strategy to improve the yield of locus associated complexes, reduce sample complexity and improve sensitivity.

Here, starting from 3 billion cells, we combined a CRISPR-dCas9 based reverse-ChIP approach with a sequential purification step targeting histones, to biochemically remove the unwanted background, prior to semi-quantitative Mass Spectrometry identification; a purification pipeline we refer to as “dCas9 targeted chromatin and histone enrichment for mass spectrometry’’ (Catchet-MS). Catchet-MS enabled specific enrichment and identification of the chromatin-bound fraction of the dCas9 bait proteins bound to the target region, while non-localized bait complexes in the cytoplasm and nucleoplasm were depleted. Exclusive selection on proteins that were identified to be differentially bound to the latent versus the transcriptionally activated HIV 5’LTR was implemented to stringently remove nonspecific background.

Using Catchet-MS as a discovery tool we identify novel and previously reported proteins preferentially associated with the latent or active HIV-1 promoter that represent potential key regulators of HIV-1 gene expression. Among these regulators, we demonstrate that the transcription factor IKZF1 is a novel transcriptional repressor of the HIV-1 promoter, required for the sequential recruitment of the polycomb repressive complex 2 (PRC2) and critical for maintenance of a repressive LTR chromatin environment characterized by the presence of the H3K27me3 chromatin mark. Remarkably, we demonstrate that the therapeutically advanced drug iberdomide (CC-220), and the FDA approved thalidomide analogues lenalidomide and pomalidomide, by targeting IKZF1 protein for CRL4^CRBN^ dependent degradation, lead to HIV-1 latency reversal in both ex vivo infected primary CD4+ T cells and cells isolated from HIV-1 infected aviremic individuals on cART. Our data suggest that modulation of IKZF1 protein expression with iberdomide, or other thalidomide derived drugs such as the FDA-approved lenalidomide and pomalidomide, represents a viable target for therapeutic intervention to reverse latency in the context of HIV-1 cure strategies.

## Results

### Catchet-MS, a target discovery strategy to identify locus-bound protein complexes at the latent and activated HIV-1 LTR in vivo

We established a CRISPR/dCas9-based reverse-ChIP proteomics strategy that incorporates a sequential purification step targeting histones which we refer to as “dCas9 targeted chromatin and histone enrichment for mass spectrometry’’ (Catchet-MS) to identify a comprehensive repertoire of protein complexes associated in vivo with the latent and active HIV promoter (**Figure 1A-H**). As a cellular model system for latency we used J-Lat 11.1, a Jurkat derived T cell line, which harbors a single latent copy of the integrated full-length HIV-1 proviral genome, defective in *env*, and in which GFP replaces the *nef* coding sequence as a reporter for viral activation (*27*) (**Figure 1A**). J-Lat 11.1 cells were first modified to stably express a multiple epitope (V5, HA, FLAG)-tagged dCas9 bait, and an 18 nucleotide single guide-RNA, targeting the hypersensitive site 2 (HS2) region of the HIV-1 promoter, downstream of the Transcription Start Site (TSS) (**Figure 1A**). We chose to target our dCas9 bait to the HS2 region downstream of Nuc-1, as recent evidence pointed to transcription elongation and RNA processing as critical steps involved in HIV-1 latency in vivo (*28*). As a negative control, we generated cells expressing HA-V5-FLAG-dCas9 and a non-targeting control gRNA (nt-gRNA). Expression of the tagged dCas9 bait construct was confirmed by Western blotting, using antibodies specifically recognizing dCas9 and the construct tags HA, V5, and FLAG (**Figure 1B**). We validated gRNA (HS2-gRNA) dependent HIV-1 promoter targeting both by a functional assay, co-transfection of J-Lat 11.1 cells together with a dCas9-VP64-p65-Rta (dCas9-VPR) fusion construct containing the VPR transcriptional activation domain (**Supplementary Figure 1A**) as well as by ChIP-qPCR analysis using a V5 epitope-based IP (**Figure 1D**). Binding of the HA-V5-FLAG-dCas9 bait is not affected by the transcriptional state of the 5’LTR as demonstrated by ChIP-qPCR in latent and PMA stimulated cells (**Figure 1E**) and does not interfere functionally with viral reactivation as demonstrated by Flow cytometry and fluorescence microscopy of GFP expression in the HA-V5-FLAG-dCas9 expressing J-Lat 11.1 cells both before and after PMA stimulation (**Figure 1C and Supplementary Figure 1B**). Thus, the targeted dCas9 bait is properly targeted to the 5’HIV LTR and does not interfere with the maintenance of latency or capacity for reactivation. From this, we infer that binding of 5’LTR associated transcription regulatory factors bound in the repressed or activated states is unaltered by dCas9 targeting.

**Figure 1.**
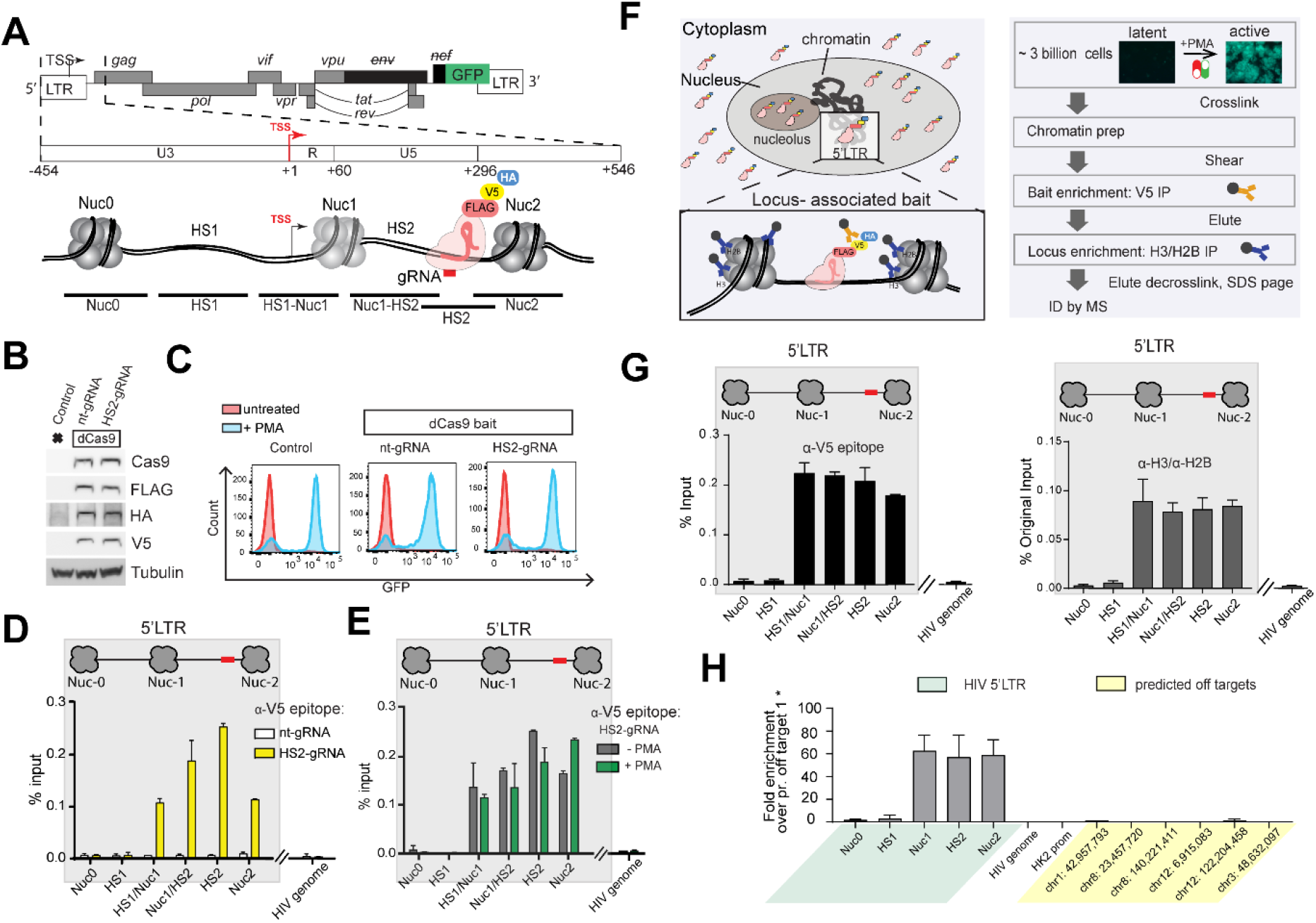
dCas9 targeted chromatin and histone enrichment for mass spectrometry (Catchet-MS), a method to isolate and identify locus-bound protein complexes in vivo. (A) Schematic representation of the genomic organization of the integrated HIV-1 provirus in J-Lat 11.1 cells, encoding GFP and containing a frameshift mutation in *env* and a partial deletion of *nef*. The 5’ LTR region is further segmented into the U3, R, and U5 regions. Three strictly positioned nucleosomes, Nuc-0, Nuc-1 and Nuc-2, delimit the nucleosome-free regions HS1 and HS2, hypersensitive to nuclease digestion, as indicated. The HS2 region, to which the multiple epitope-tagged HA-V5-FLAG-dCas9 bait is guided, is indicated. The amplicons used to scan the chromatin region in ChIP-qPCR are shown. (B) Western blot analysis indicates expression of multiple epitope-tagged HA-V5-FLAG-dCas9 bait in modified J-Lat 11.1 cells using antibodies specific for Cas9, V5, FLAG and HA as indicated. Parental J-Lat 11.1 cell lysate is used as a negative control and α-Tubulin is used as a loading control. (C) Flow cytometry histograms show the distribution of GFP positive cells in unstimulated and PMA stimulated control J-Lat 11.1 cells, cells expressing the dCas9 bait and a non-targeting gRNA (nt-gRNA) and cells expressing the bait and the HS2 targeting gRNA (HS2-gRNA). (D) ChIP-qPCR analysis with anti V5 epitope affinity beads indicate specific enrichment of the HA-V5-FLAG-dCas9 bait over the guided HIV-1 LTR region. White bars represent data generated in cells expressing the dCas9 bait together with a non-targeting gRNA (nt-gRNA), yellow bars represent data generated in cells expressing the bait and the single guide RNA targeting the HS2 region of the HIV-1 5’LTR (HS2-gRNA). Data show a representative experiment, error bars represent the standard deviation (±SD) of two separate real-time PCR measurements. HIV-1 5’LTR sequences recovery is calculated as a percentage of the input. (E) ChIP-qPCR analysis with anti V5 epitope affinity beads in latent (-PMA; grey bars) and PMA treated cells (+PMA; green bars) expressing the dCas9 bait and the HS2-gRNA. Data are the mean of 2 independent experiments (±SD). HIV-1 5’LTR sequences recovery is calculated as a percentage of the input. (F) Schematic representation of the dCas9 bait cellular localization (left panel) and the Catchet-MS workflow (right panel). Approximately 3 billion cells per condition are cross-linked with formaldehyde to stabilize the protein-protein and protein DNA interaction. Following a stringent chromatin enrichment protocol, the cross-linked chromatin is isolated and fragmented by ultrasound sonication. The dCas9 containing complexes are immunoprecipitated using anti V5 antibody conjugated affinity beads, eluted from the beads, and used as input material for a second round of purification with anti-histones (H3, H2B) antibody-conjugated beads, in order to remove the non-chromatin bound fraction of the HA-V5-FLAG-dCas9 bait complexes and to enrich for the locus associated bait complexes. Immunoprecipitated material is finally decrosslinked, resolved on an SDS-page and prepared for mass spectrometry analysis. (G) (Left panel) ChIP-qPCR analysis with anti V5 epitope affinity beads in chromatin fraction prepared from unstimulated cells indicates specific enrichment of HA-V5-FLAG-dCas9 over the HIV-1 5’LTR. Data show a representative experiment, error bars represent the standard deviation (SD) of two separate real-time PCR measurements, HIV-1 5’LTR levels are calculated as percentages of the input. The V5 affinity-purified chromatin (represented in top panel) was eluted and used as input for a sequential immunoprecipitation with a mix of histone H2B and H3 conjugated affinity beads and the isolated material was analyzed by qPCR (Right panel). ChIP-qPCR with anti H3/H2B conjugated affinity beads indicates 5’LTR enrichment of HA-V5-FLAG-dCas9 bait after sequential V5/histone affinity purification.Data show a representative experiment, error bars represent the standard deviation (SD) of two separate real-time PCR measurements, HIV-1 5’LTR levels in both the V5 and histone affinity purification are calculated as percentages of the same original input. (H) Average coverage profiles using ChIP sequencing reads mapped 500bp upstream and downstream of the peak center at the 5’LTR region of the HIV genome (’Targets’) to the respective coverage around the predicted off targets (’OffTargets’). ’Start’ denotes the starting base pair of the aforementioned 1kb region around the peak centers and ’End’ the ending base pair respectively.

The optimal HA-V5-FLAG-dCas9 bait IP enrichment strategy was determined by examining specific enrichment of the locus using antibodies directed against the HA, V5, and FLAG epitopes in ChIP-qPCR experiments (**Supplementary Figure 1C-D**). While antibodies directed against each tag resulted in efficient and specific enrichment of the targeted HIV-1 DNA region, we chose to use anti V5 affinity beads targeting the V5 epitope as the first step in the large-scale locus enrichment strategy as it demonstrated both good purification yield as well as the highest signal to noise ratio (**Supplementary Figure 1D**).

The large number of non-localized HA-V5-FLAG-dCas9 bait molecules e.g. accumulating in the cytoplasmic fraction (**Figure 1F left panel, Supplementary Figure 2A**) or in the RNA-rich nucleolar compartment (*29, 30*) presented a caveat in our single locus chromatin proteomics reverse-ChIP approach; the non-targeted dCas9 bait molecules and associated proteins are also captured through the purification pipeline thus interfering with reliable identification of the locus associated bait interactome. To eliminate this source of background we modified our in-house ChIP protocol and included a Chromatin enrichment step (adapted from the Chromatin enrichment for proteomics (CheP) protocol (*31*)) as well as a histone antibody based enrichment step into our purification pipeline (**Figure 1F right panel**). Implementation of these steps resulted in reduced recovery of HA-V5-FLAG-dCas9 (due to loss of bait-associated complexes not associated with histones; **Supplementary Figure 2B**) while the HIV-1 promoter was still efficiently recovered (**Figure 1G**). The number of proteins identified was also reduced in consecutive steps of the pipeline (**Supplementary Figure 2C**) while proteins associated with the chromatin compartment were enriched (**Supplementary Figure 2D**). Conversely, proteins associated with the nucleolus (**Supplementary Figure 2E**) or the cytoplasm and cell membrane (**Supplementary Figure 2D**) were depleted.

We subjected the genomic DNA recovered after the Catchet-MS pipeline to next generation sequencing and qPCR to determine the specificity of the approach. Both in ChIP-seq (**Supplementary Figure 2F and G**) and its confirmation by ChIP qPCR (**Figure 1H**) we observed a strong enrichment of the HIV-1 5’LTR while enrichment at human chromosomes, aside from centromeric artifacts common to ChIP-seq and a few minor off-target peaks (**Supplementary Table 1**), is low. Zooming in on predicted off-target regions (http://www.rgenome.net/cas-offinder/) in the NGS-data (**Supplementary Figure 2G**) and confirmation by qPCR (**Figure 1H**) demonstrates the absence of traceable peaks or any substantial PCR signal for these chromosomal loci.

In summary our results demonstrate that Catchet-MS enables highly specific enrichment of the targeted HIV-1 5’LTR bound proteome while cellular proteins are depleted, indicating its suitability as a discovery tool to identify HIV-1 5’LTR associated host factors.

### Cachet-MS identifies known and novel host factors associated with the latent and active HIV-1 promoter

We scaled up the Catchet-MS workflow (**Figure 1F**), to approximately 3 billion cells as starting material in the unstimulated and PMA stimulated conditions to model J-Lat 11.1 HIV-1 latency and activation. **Figure 2A** presents a heatmap of the mass spectrometry analysis performed at different steps of the Catchet-MS purification pipeline. Sample complexity is high in the input samples and are characterized by a large number of high protein intensity values **Figure 2A**. As expected, the abundance of the HA-V5-FLAG-dCas9 bait is low, in these input samples (**Figure 2B and C**). Anti-V5 affinity purification results in efficient enrichment for the HA-V5-FLAG-dCas9 bait-containing complexes and a marked reduction in sample complexity (**Figure 2A-C**). Finally, re-immunoprecipitation using anti-Histone H2B and H3 conjugated affinity beads leads to a clear difference in protein abundances between the unstimulated and PMA stimulated samples, as shown in the last two lanes of **Figure 2A**.

**Figure 2.**
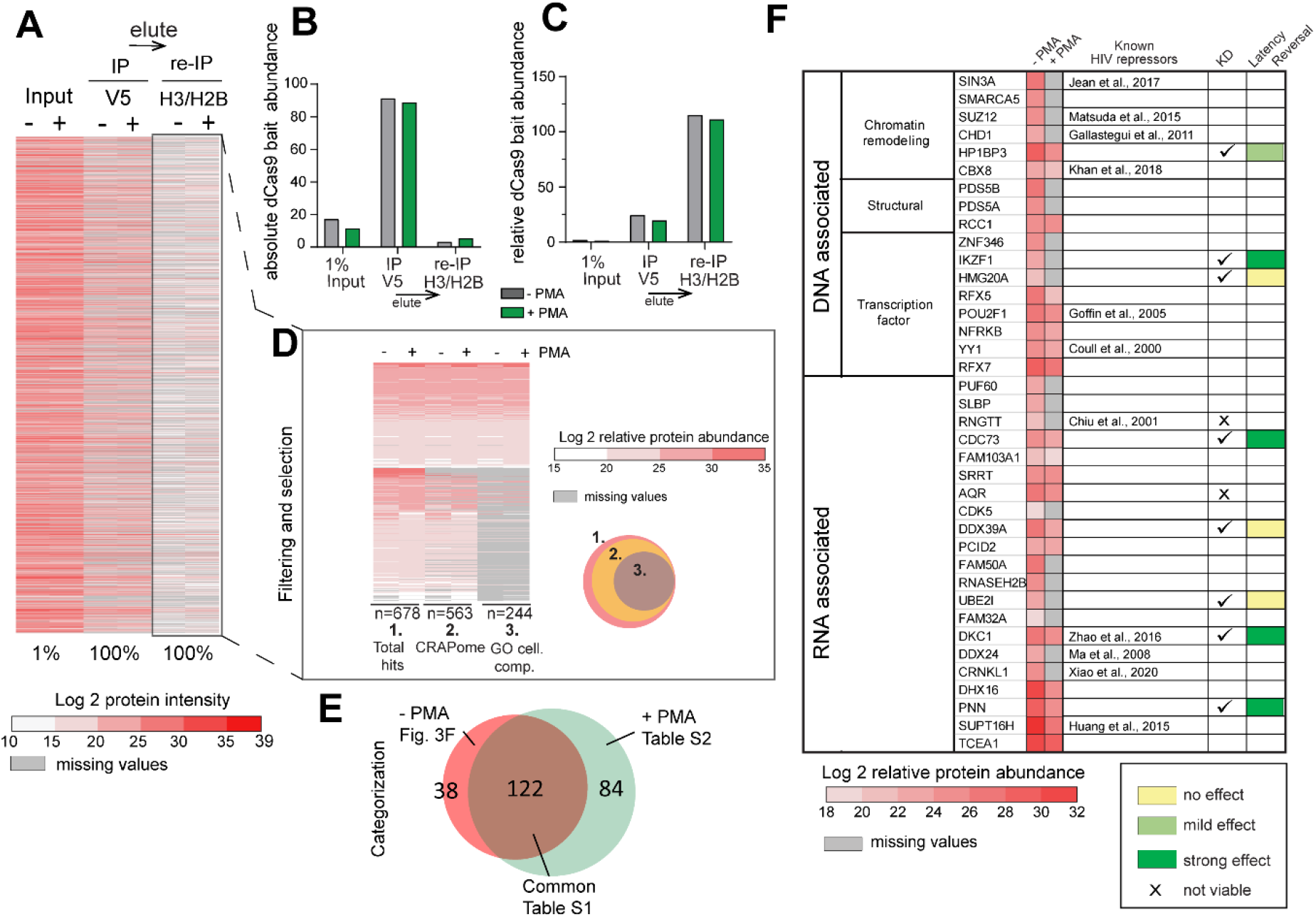
**Cachet-MS identifies known and novel host factors associated with the latent and active HIV-1 promoter**(A) Heatmap displays the protein content at each step of the Catchet-MS purification pipeline. The colors represents the Log2 transformation of the protein intensities scores. Values corresponding to the V5 based immunopurification are adjusted to 100% as only a fraction (1:40) of the material that went into the second, histone based (H2B/H3) immunopurification, was analyzed by mass spectrometry. For the V5/histone experiment, 100% of the material was subjected to mass spectrometry. Missing values are represented by grey lines. The heatmap represents data from one Catchet-MS experiment, input material for the experiment corresponds to chromatin generated from a starting material of 3 billion cells per condition. (B) Bar plots showing the absolute HA-V5-FLAG-dCas9 bait abundance at each step of the purification pipeline. The absolute abundance was calculated based on protein/peptide spectral intensity values and adjusted to the fraction of material analyzed by mass spectrometry. The values are displayed in arbitrary units. (C) Bar plots showing the relative HA-V5-FLAG-dCas9 bait abundance at each step of the purification pipeline. The relative abundance was calculated based on protein/peptide spectral intensity values and normalized to the total protein content. The values are displayed in arbitrary units. (D) Filtering criteria applied to Catchet-MS data. Hits are filtered by comparison with the Crapome repository for contaminants of affinity purification experiments . as well as through GO categorization to select hits present in the cell nucleus and classified to be RNA bound, DNA bound or alternatively bound to histones, transcription factors or associated to chromatin. Colors represent the Log2 transformation of the protein relative abundance. The protein relative abundance was calculated based on protein/peptide spectral intensity values and normalized to the total protein content. Missing values are represented by grey lines. (E) Venn diagram graphically summarizing the overlap between the hits identified to be exclusively, or more abundantly, associated with the unstimulated state (-PMA) and the hits associated with the activated state (+PMA). (F) Selection and functional classification of hits associated with the unstimulated state and considered for a potential role in the maintenance of HIV-1 latency. The first columns of the table summarizes in a heatmap the Log2 trasfomation of the protein relative abundance values in the unstimulated (–PMA) sample and the activated (+PMA) sample. Missing values are represented by grey lines. Candidates previously reported to restrict HIV-1 expression are referenced in the third column. Checkmarks indicate the hits functionally validated by shRNA mediated depletion experiments (Supplementary Figure 3C). Colored boxes (yellow to dark green) summarize the effect of the target depletion on HIV-1 expression and refers to the effects summarized in S3.

Analysis of the MS data from the final locus enrichment step resulted in a list of 678 proteins, identified with high confidence, Mascot score > 100. We then applied stringent and unbiased selection criteria, Illustrated in **Figure 3D**, to narrow down the list of candidates by removing potential contaminants. First, we filtered out frequent contaminants using the Crapome database (https://www.crapome.org/), second we applied a GO cellular compartment and function analysis (GO-terms nucleus, chromatin-associated, histones bound, bound to transcription factors, DNA associated, or RNA associated) to include only nuclear proteins in our further analysis. Finally, we determined the differential proteome of latent versus activated HIV-1 LTR bound factors, by excluding from further analysis proteins common to both conditions, which include potential non-specific background proteins. While this may have also resulted in elimination of potentially functionally relevant factors bound to both transcriptional states of the promoter, it allowed for a stringent removal of potential background. Applying these criteria, we compiled a list (n=38) of proteins predominantly associated with the HIV-1 promoter in the latent state (**Figure 2E and F**), and a list (n=84) of factors predominantly associated with the activated HIV-1 promoter following treatment with PMA (**Figure 2E and Supplementary Table 2**) while (n=122) common proteins, including the dCas9 bait and potential contaminants, were found associated with the HIV-1 5’LTR under both conditions (**Figure 2E and Supplementary Table 3**)

**Figure 3.**
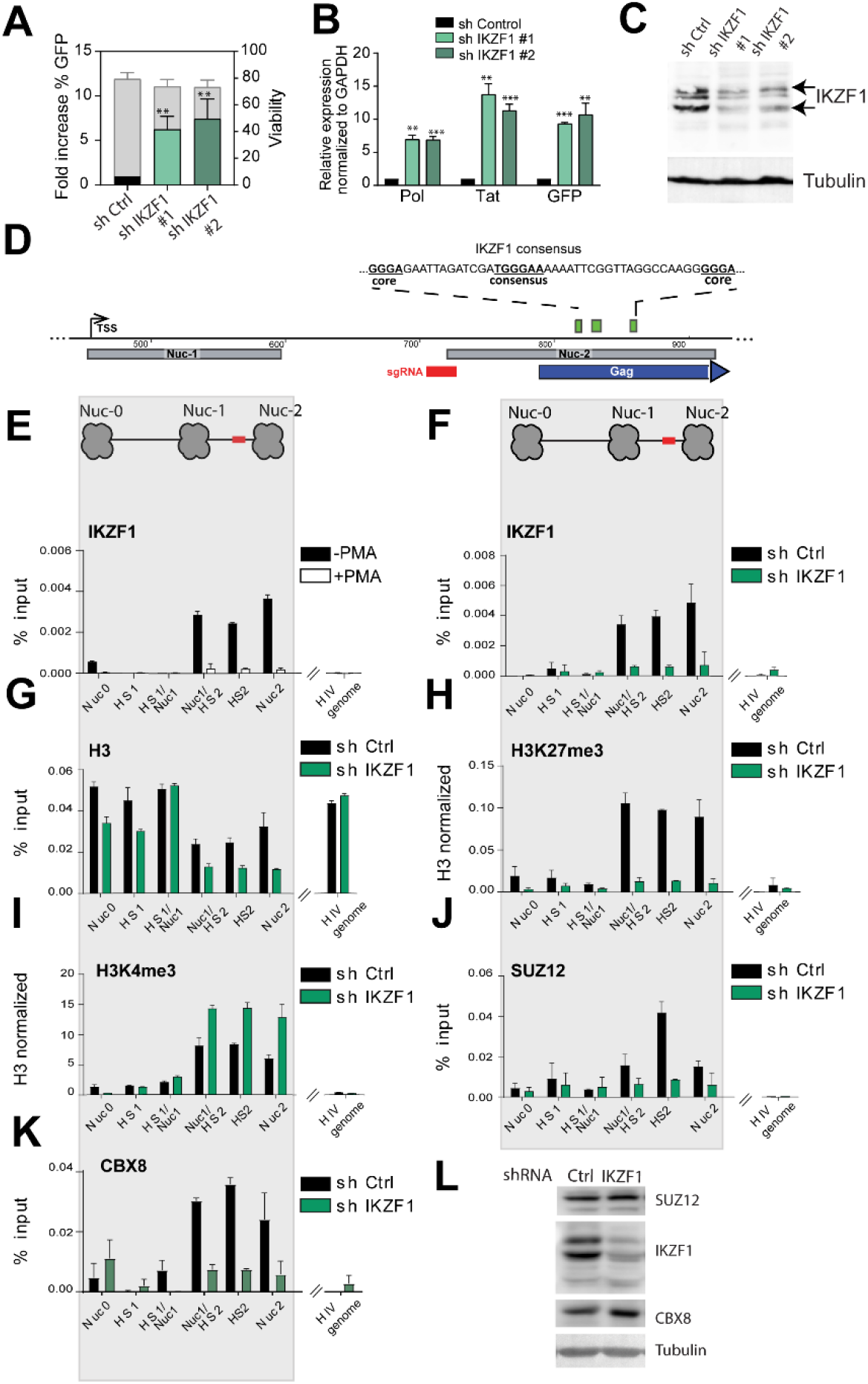
IKZF1, required for maintenance of HIV-1 latency, binds downstream of the latent HIV-1 5’LTR to promote PRC2/PRC1 recruitment and establish a repressive chromatin environment. (A) Bar plot showing the fold increase in % GFP positive cells (left y axes) measured by FACS analysis, following IKZF1 depletion in J-Lat 11.1 with two different shRNA constructs (#1 and #2). Data are the mean of three independent experiments (±SD). The right y axis represents the percentage of live cells. (B) qRT-PCR analysis measuring expression of HIV genes (pol, GFP, tat) in J-lat 11 transduced with scramble shRNA (sh Control) and sh IKZF1 #1 and #2. Data, normalized to GAPDH are represented as fold enrichment over sh Control and are the mean of three independent experiments (±SEM). Statistical significance was calculated using a ratio paired t test, * – p<0,05; **– p<0,01– ***p<0,001. (C) Western blotting for IKZF1 in J-lat 11 transduced with scramble shRNA (sh Control) and shIKZF1 #1 and #2 iondicates depletion of IKZF1. α-Tubulin is used as a loading control. (D) Putative IKZF1 binding site (835-840) downstream of the HIV-1 5’-LTR in the proximity of the sequence targeted by the gRNA. The IKZF1 binding site is composed of a consensus sequence (TGGGAA/T) and at least one more extra core site (GGGA) in a 40 bp range (Li et al., 2011). (E) ChIP-qPCR analysis with IKZF1 antibody in latent cells and PMA stimulated J-Lat 11.1 cells as indicated. (F) ChIP qPCR analysis with IKZF1 antibody in J-Lat 11. 1 cells transduced with scramble shRNA (shControl) and shIKZF1. Data in (E) and (F) are presented as % input, error bars represent the standard deviation (SD) of two separate real-time PCR measurements. (G, H, I) ChIP-qPCR using antibodies specific for distinct histone marks in J-Lat 11. 1 cells transduced with scramble shRNA (shControl) and shIKZF1. Total histone H3 (G), H3K27me3 (H), H3K4me3 (I). Total histone H3 data (G) are represented as % input mean (±SD), histone marks data (H)(I) are expressed as fold change over H3 signal (±SD). (J) ChIP-qPCR analysis with SUZ12 in J-Lat 11. 1 cells transduced with scramble shRNA (shControl) and shIKZF1. (K) ChIP-qPCR analysis with CBX8 in J-Lat 11. 1 cells transduced with scramble shRNA (shControl) and shIKZF1. Data in (J) and (K) are presented as % input, error bars represent the standard deviation (SD) of two separate real-time PCR measurements. The ChIP analysis presented (E-K) are a representative experiment, biological replicate experiments are shown in S8. (L) Western blotting for SUZ12, IKZF1, CBX in J-lat 11 transduced with scramble shRNA (sh Control) and shIKZF1. α-Tubulin is used as a loading control.

## Functional validation of proteins identified to associate with the latent promoter

In search of putative molecular targets for therapeutic inhibition and HIV-1 latency reversal, we exclusively selected protein hits that were distinctly associated with the latent HIV-1 LTR, in vivo, comprising 38 factors presented in the table in **Figure 2F**. Of these latent HIV-1 LTR-associated candidates, a significant number (n=11) were previously reported to restrict viral expression (**Figure 2F** referenced in the table) and serve as positive controls that validate the quality of the experimental approach, providing confidence towards the veracity of the candidate list. We evaluated the effect of shRNA-mediated knockdown of a selection of the novel factors we identified, on the HIV-1 5’LTR-driven GFP reporter by flow cytometry and on the expression of HIV-1 genes Gag, pol, and Tat by RT-PCR in J-Lat 11.1 cells (**Figure 2F; Supplementary Figure 3**). Depletion of a number of the selected candidates (n=5/10, namely HP1BP3, IKZF1, CDC73, DKC1, and PNN), resulted in significant (p <0.05) upregulation of HIV-1 genes and an increase in the percentage of GFP positive cells (green boxes, **Figure 2F** and **Supplementary figure 3**), while for a small number of factors tested (N=3/10), we did not observe a significant change in HIV-1 expression (yellow boxes, **Figure 2F**) and the remaining factors were found to be essential for cell viability (x mark, **Figure 2F**). Our data suggested that the selected n=5 candidates are required for repression of HIV-1 expression in J-Lat 11.1 cells.

### IKZF1 binds to the latent HIV-1 promoter and is repressive to HIV-1 transcription

Catchet-MS identified the IKAROS Family Zinc Finger 1 protein (IKZF1) as one of the factors to be exclusively associated with the repressed HIV-1 promoter (**Figure 2F**). IKZF1, a critical factor required for normal T cell development that can act both as a transcriptional repressor and activator (*32–34*), has been reported to regulate gene expression through its association with the nucleosome remodeling and deacetylase complex (NuRD) and the positive transcription elongation factor (P-TEFb) (*34–38*). Additionally, IKZF1-mediated gene silencing in T cells has been shown to be facilitated by its interaction with the Polycomb repressive complex 2 (PRC2), which promotes histone H3 lysine 27 trimethylation, a mark of transcriptionally silent chromatin (*39*). Consistent with its role as a repressive factor in T cells, depletion of IKZF1 in our functional testing screen using independent shRNA clones led to the strongest reactivation of HIV-1 expression (5 to 10 fold), as measured by the increase of GFP-positive cells in flow cytometry and qRT-PCR for HIV-1 genes (**Figure 3A-C**). To ensure that the observed latency reversal is not subject to clonal effects in J-Lat 11.1 cells, we depleted IKZF1 in J-Lat A2 cells, harboring an integrated HIV 5’ LTR-driven GFP reporter and observed similar reversal of HIV-1 latency (**Supplementary Figure 4A-B**).

The IKZF family consists of several family members (*37, 40*), although IKZF1 is the only family member detected in our mass spectrometry data. We assessed the expression of the other IKZF family members in Jurkat cells (RNA sequencing (*41*)) and proceeded with shRNA mediated depletion of IKZF2 and 5, which are expressed in Jurkat cells (**Supplementary Figure 5A**). Depletion of IKZF2 and IKZF5 also resulted in HIV-1 latency reversal, although less pronounced compared to IKZF1 (**Supplementary Figure 5B**). These results are consistent with the lower levels of expression of IKZF2 and IKZF5 in T cells, compared to IKZF1, and with the well-established heterodimeric interactions among IKZF family members (*42–44*). We next sought to confirm and examine the association of IKZF1 with the latent HIV-1 LTR. Downstream of the 5’LTR, at position +818/+864 we identified a putative IKZF binding site, which is composed of a consensus TGGGAA/T sequence and two extra GGGA core sites (*45*), (**Figure 3D**). To assess IKZF1 binding at the predicted sites, we performed ChIP-qPCR experiments in latent and PMA treated J-Lat 11.1 cells using antibodies specific for IKZF1 (**Figure 3E**). In agreement with the Catchet-MS data, ChIP experiments in latent J-Lat 11.1 cells demonstrated IKZF1 enrichment between HIV-1 HS2 and Nuc2, (**Figure 3E**). Importantly, activation of HIV-1 transcription by PMA treatment abrogated IKZF1 binding (**Figure 3E and Supplementary Figure 6A**), consistent with the notion that IKZF1 may be important for maintenance of HIV-1 transcriptional repression. To confirm the specificity of IKZF1 binding, we also performed ChIP in control J-Lat 11.1 cells, infected with a scramble shRNA vector (Sh Control), and in cells infected with an shRNA vector targeting the IKZF1 mRNA (sh IKZF1). Our data show clear enrichment of IKZF1 binding over its predicted binding region in the sh Control cell line together with a dramatic loss of binding upon shRNA mediated depletion in the IKZF1 depleted cells (**Figure 3F and Supplementary Figure 6B**), thus confirming the specificity of the antibody. To further control for the specificity of the IKZF1 signal we also monitored enrichment of IKZF1 at endogenous IKZF1 targets previously reported to be enriched for IKZF1 binding in B cells (*46*) (**Supplementary Figure 6C**). We then examined the effect of IKZF1 depletion on the chromatin state of the region by histone ChIP-qPCR experiments using antibodies against the active chromatin modification H3 lysine 4 tri-methylation (H3K4me3), the repressive chromatin modifications H3 lysine 27 tri-methylation (H3K27me3) and H3 lysine 9 tri-methylation (H3K9me3), using the total histone H3 signal for normalization (**Figure 3G-I; Supplementary Figure 7A-D**).

Upon IKZF1 depletion the chromatin state is characterized by a drastic loss of enrichment for the H3K27me3 mark (**Figure 3H; Supplementary Figure 7B**), a moderate increase of the H3K4me3 mark (**Figure 3I; Supplementary Figure 7C**) and a concomitant decrease in the repressive chromatin mark H3K9me3 (**Supplementary Figure 7D**). Similar data were obtained upon IKZF1 depletion in J-Lat A2 cells (**Supplementary Figure 7E-I**), thus confirming that the changes in chromatin states are not subject to clonal effects.

We also assessed the histone modification profile in latent and PMA stimulated J-lat 11.1 cells (**Supplementary Figure 8A-D**). Mimicking what was observed upon IKZF1 depletion, treatment with PMA led to an increase in H3K4 tri-methylation (**Supplementary Figure 8B**) and a concomitant decrease in the repressive chromatin marks H3K27me3 (**Supplementary Figure 8C**) and H3K9me3 (**Supplementary Figure 8D**). IKZF1 depletion, however, led to a more prominent loss of enrichment for the H3K27me3 mark (**Figure 3H**).

In T cells, IKZF1 is necessary for mediating gene silencing through the recruitment of the Polycomb repressive complex 2 (PRC2) and the deposition of the H3K27me3 mark (*39*). Interestingly, our Catchet-MS experiment identified SUZ12, a core subunit of the PRC2 complex, and CBX8, a subunit of the Polycomb repressive complex 1 (PRC1) that acts as a reader of the H3K27me3 modification (*47–49*) to be also associated with the HIV-1 5’LTR in its latent state. We therefore conducted ChIP experiments to probe for the IKZF-1 mediated binding of SUZ12 to the region to verify the hypothesis that IKZF1 may be necessary for recruitment of PRC2 complex to the HIV-1 5’LTR. We additionally examined the recruitment of CBX8 to check whether the reduction in H3K27me3 mark deposition observed in absence of IKZF1 affects CBX8 binding. Consistent with this hypothesis, our data shows that depletion of IKZF1 leads to reduced enrichment of SUZ12 (**Figure 3J and Supplementary Figure 8E**) and CBX8 (**Figure 3K**) over the HS2 5’LTR region. Importantly, as shown by western blotting, depletion of IKZF1 does not lead to decreased levels of expression of SUZ12 and CBX8, suggesting that the effects observed are exclusively consequence of the absence of IKZF1 recruitment to the region (**Figure 3L**).

### Targeting IKZF1 by iberdomide treatment reverses HIV-1 latency in ex vivo infected primary CD4+ T cells

Thalidomide-derived immunomodulatory drugs (IMIDs), have been shown to cause selective ubiquitination and degradation of IKZF1 and its related family member IKZF3 (*50, 51*). The mechanism of action for IMID activity lies in their affinity for cereblon (CRBN), which is part of the cullin-ring finger ligase-4 cereblon (CRL4^CRBN^) E3 ubiquitin ligase complex. Iberdomide is a novel compound with higher affinity for CRBN and the lowest IC50 of all thalidomide analogues, and is currently under clinical development for the treatment of systemic lupus erythematosus (SLE) and relapsed/refractory multiple myeloma (RRMM).

Binding of iberdomide to CRBN modulates the E3 ubiquitin ligase activity of the complex, increasing its affinity for IKZF1 and IKZF3, leading to their ubiquitination and proteasome-dependent degradation. Therefore, we set out to establish if iberdomide treatment in latent HIV-1 infected cells would induce degradation of IKZF1 and lead to HIV-1 latency reversal.

As the main reservoir of latent HIV-1 in infected individuals comprises resting CD4+ T cells, we ex vivo infected, without prior activation, CD4+ T cells obtained from healthy donors with a defective full-length HIV-1 virus harboring a luciferase reporter to establish latent infections (**Supplementary Figure 9A**) (*52*). Treatment of latent HIV-1-infected primary CD4+ T cells with iberdomide for 48 hours resulted in moderate but significant (p < 0.05) reversal of latency, as observed by an increase in the mean luciferase activity compared to the untreated control (**Figure 4A**) in all 6 donor CD4+ T cell lines examined. Bromodomain end extra-terminal domain (BET) inhibitors have been reported to act synergistically with IMIDs in treatment of refractory forms of multiple myeloma(*53, 54*). Therefore, we examined whether these compounds also synergize in the context of HIV-1 latency reversal. Indeed, co-treatment with Iberdomide and JQ1 resulted in a robust and synergistic HIV-1 latency reversal (**Figure 4A**). To confirm that treatment with iberdomide results in degradation of IKZF1 at the protein level, we performed western blot analysis of primary cells treated for 48 hours with iberdomide. As expected, treatment with iberdomide, alone or in combination with JQ1, resulted in degradation of IKZF1, as shown in **Supplementary Figure 9B**. Importantly, treatment with iberdomide did not affect IKZF1 expression at the level of transcription (**Supplementary Figure 9C**), consistent with the known post-transcriptional mechanism by which IMIDs target IKZF1 for CRBN mediated degradation. Conversely, mRNA levels of the endogenous IKZF1 targets p21 and cMyc (*55, 56*) were significantly affected by the treatment (**Figure 9C**).

**Figure 4.**
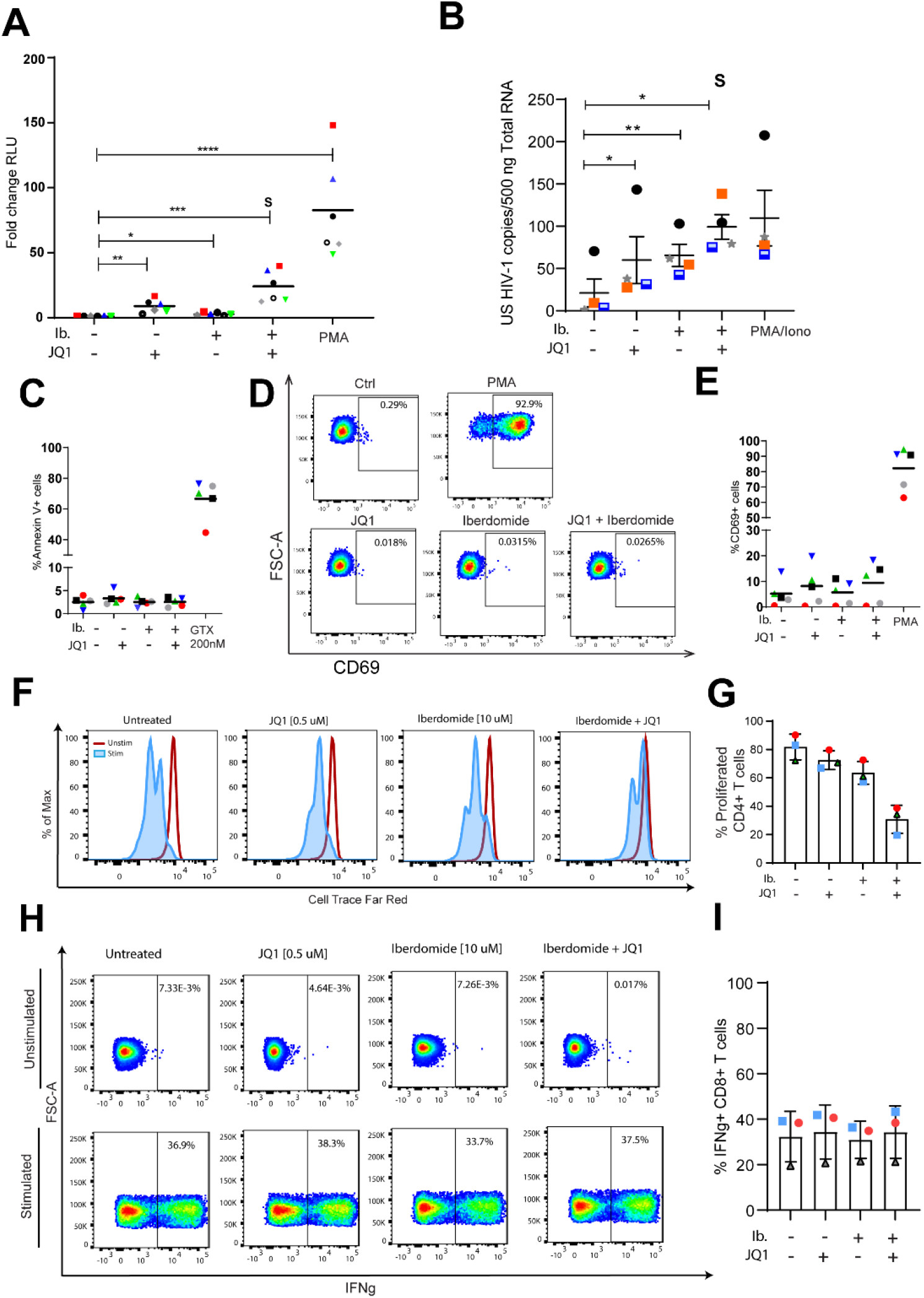
Targeting Targeting IKZF1 by iberdomide treatment reverses latency in primary CD4+ T cells obtained from cART suppressed HIV-1 infected patients with minimal effect on toxicity and effector function. (A) The latency reversal activity of iberdomide alone (10uM) and in combination with JQ1 (500nM) was tested, in primary human CD4+ T cells ex vivo infected to harbour latent HIV-1. The dot plot in panel A shows the fold increase in luciferase activity after treatment as indicated. Each dot represents a single measurement while the black horizontal lines represent the average fold increase for each treatment in the pool of donors. Experiments were performed in duplicate using cells obtained from 6 healthy donors. S represents synergism, calculated by using the coefficient of drug interaction (CDI) method. Statistical significance was determined by repeated measures one-way ANOVA on the log-transformed fold changes followed by Tukey’s multiple comparison test * – p < 0,05, ** – p < 0,01, *** – p < 0,001, ****– p < 0,0001. (B) Graph panel showing the average levels of unspliced cell-associated HIV-1 RNA in CD4+ T cells isolated from five HIV-1 infected, aviremic participants upon treatment with Iberdomide, JQ-1 or combination as measured by nested qPCR. PMA/Ionomycin was used as a positive control. CD4+ T cells were isolated from PBMCs from HIV-1 infected donors and treated as indicated for 24 hours. Statistical significance was calculated using paired two-tailed t-test, * – p < 0,05, ** – p < 0,01. (C) Percentage of cells expressing the Annexin V marker of apoptosis in primary CD4+ T cells treated with iberdomide, JQ1 and the combination of both compounds for 24 hours. Treatment with a toxic concentration of Gliotoxin (GTX) 200nM was used as a positive control. Experiments were performed in uninfected cells obtained from five healthy donors, represented by the dots. (D) Representative flow cytometry plot of extracellular CD69 marker staining analysis in primary CD4+ T cells, upon treatment with JQ1, iberdomide, or a combination of both compunds for 24 hours. CD69 expression was assessed by extracellular staining and analyzed by flow cytometry. (E) Percentage of cells expressing the CD69 marker of cell activation in primary CD4+ T cells from 5 healthy donors as described in E. Treatment with PMA was used as a positive control. (F)Representantive histogram of proliferative capacity of unstimulated or aCD3/CD28 stimulated CD4+ T cells in the presence or absence of LRAs. Cells were stained with a proliferation dye and analyzed 72 hours later by flow cytometry. Dividing cells show decreased intensity of prolifertion dye as it becomes diluted upon cell division. (G) Percentage of proliferated CD4+ T cells from 3 healthy donors as described in G. (H) Representative flow cytometry plots of IFNg production analysis in unstimulated and stimulated primary CD8+ T cells after treatment with LRAs. Cells were treated as indicated for 18 hours followed by PMA/Ionomycin (50 ng / 1 µM) stimulation for 7 hours in the presence of a protein transport inhibitor or remained unstimulated. IFN-g production was assessed by intracellular staining and analyzed by flow cytometry. Numbers in the plot show percentage of IL-2 producing cells. (I) Percentage of INFg producing CD8+ T cells from 3 healthy donors as described in H.

To validate our findings in a more relevant setting, we tested the efficacy of Iberdomide in CD4+ T cells obtained from virologically suppressed HIV-1 patients. Treatment with iberdomide alone resulted in a significant increase in cell-associated HIV-1 unspliced RNA in all 4 donors compared to control, and was comparable to that observed after treatment of cells with the BET inhibitor JQ1 (**Figure 4B**). Remarkably, in cells derived from HIV-1 infected, aviremic participants we also observe a robust, synergistic increase in cell-associated HIV-1 RNA upon co-treatment with Iberdomide and JQ-1 (**Figure 4B**). As observed in primary CD4+ T cells, treatment of Iberdomide alone of in combination with JQ-1 in cells derived from aviremic HIV infected participants led to changes in gene expression of the IKZF1 targets p21 and c-Myc (**Suppelementary Figure 9D**).

Confounding factors for potential clinical use of candidate LRAs is toxicity and the possibility of inducing unwanted global immune activation, thus leading to serious adverse effects. Potential toxicity and immune activation caused by iberdomide treatment in primary CD4+ T cells was determined by Annexin V and CD69 staining, followed by flow-cytometry. CD4+T cell viability, measured by Annexin V staining, following iberdomide treatment alone or in combination with JQ-1 was not significantly affected (**Figure 4C; Supplementary Figure 10A-C**); neither did the treatment cause global T cell activation, measured by extracellular CD69 staining (**Figure 4D-E; Supplementary Figure 10B-C**).

HIV reactivation strategies that aim for a cure will require modulation or boosting of the immune compartment to promote viral clearance. Given the known immunomodulatory role of Iberdomide and Thalidomide-derived drugs, we investigated the effects of Iberdomide treatment alone or in combination with JQ-1 on T cell proliferation and functionality. As previously reported in the literature, and consistent with their anti-lymphoproliferative capacity (*57*), we observed decreased proliferation upon treatment of CD4+ and CD8+ primary T cells (**Figure 4F and G**; **Supplementary Figure 10D and E**). Lastly, we assessed the effect of iberdomide on T cell functionality, by measuring cytokines IFNγ and IL-2 in CD4+ and CD8+ primary T cells. Iberdomide treatment alone or in combination with JQ-1 does not hamper the production of cytokines IFNγ (**Figure 4H and I; Supplementary Figure 11A and B**) and IL-2 (**Supplementary Figure 11C-F**) upon stimulation of CD4+ or CD8+ T cells and, therefore does not diminish T cell functionality. On the contrary, and consistent with the literature (*58*), we observe an increase in the production of IL-2 upon treatment with the LRAs (**Supplementary Figure 11C-F**).

### Targeting IKZF1 by FDA-approved thalidomide analogues reverses latency in cells obtained from cART suppressed HIV-1 infected patients

Although clinically advanced, iberdomide is yet to be approved by drug administrations and, therefore, cannot be presently applied to the clinic. Other thalidomide analogues, namely lenalidomide and pomalidomide, are FDA-approved drugs employed in the treatment of relapsed and refractory multiple myeloma (MM), leading to improved patient survival rates. In addition, pomalidomide is also approved for the treatment of AIDS-related Kaposi sarcoma and has proven to be well tolerated in people living with HIV, without causing drug-drug interactions with ART (*59*). These drugs have thus high potential for translational studies and direct use in the clinic, and are worth investigating in the context of HIV-1 latency reversal and cure studies.

In order to characterize the role of FDA-approved pomalidomide and lenalidomide in HIV-1 latency reversal we assessed their efficacy in ex vivo treated primary CD4+ T cells obtained from aviremic HIV-infected donors (**Table 1**). Cells were treated with different concentrations of pomalidomide and lenalidomide for 18-24 hours and latency reversal was analyzed both by FISH-Flow using probes against the viral RNA (vRNA) that target the *gag/pol* region and CA HIV-1 RNA targeting the US HIV-1 RNA. Our results show that both pomalidomide and lenalidomide are able to significantly reverse HIV-1 latency in ex vivo treated CD4+ T cells from people living with HIV (PLWH) as observed in increased vRNA+ cells upon treatment (**Figure 5A and B; Supplementary Figure 12A**). Besides, we show that bulk US HIV-1 RNA is also significantly upregulated in CD4+ T cells from PLWH upon treatment with both pomalidomide (**Figure 5C and D**) and lenalidomide (**Figure 5E and F**). Besides, we confirmed that treatment with Pomalidomide and Lenalidomide leads to degradation of IKZF1 protein in CD4+ T cells by western blot after 24 hours of treatment (**Supplementary Figure 12B-C**).

**Figure 5.**
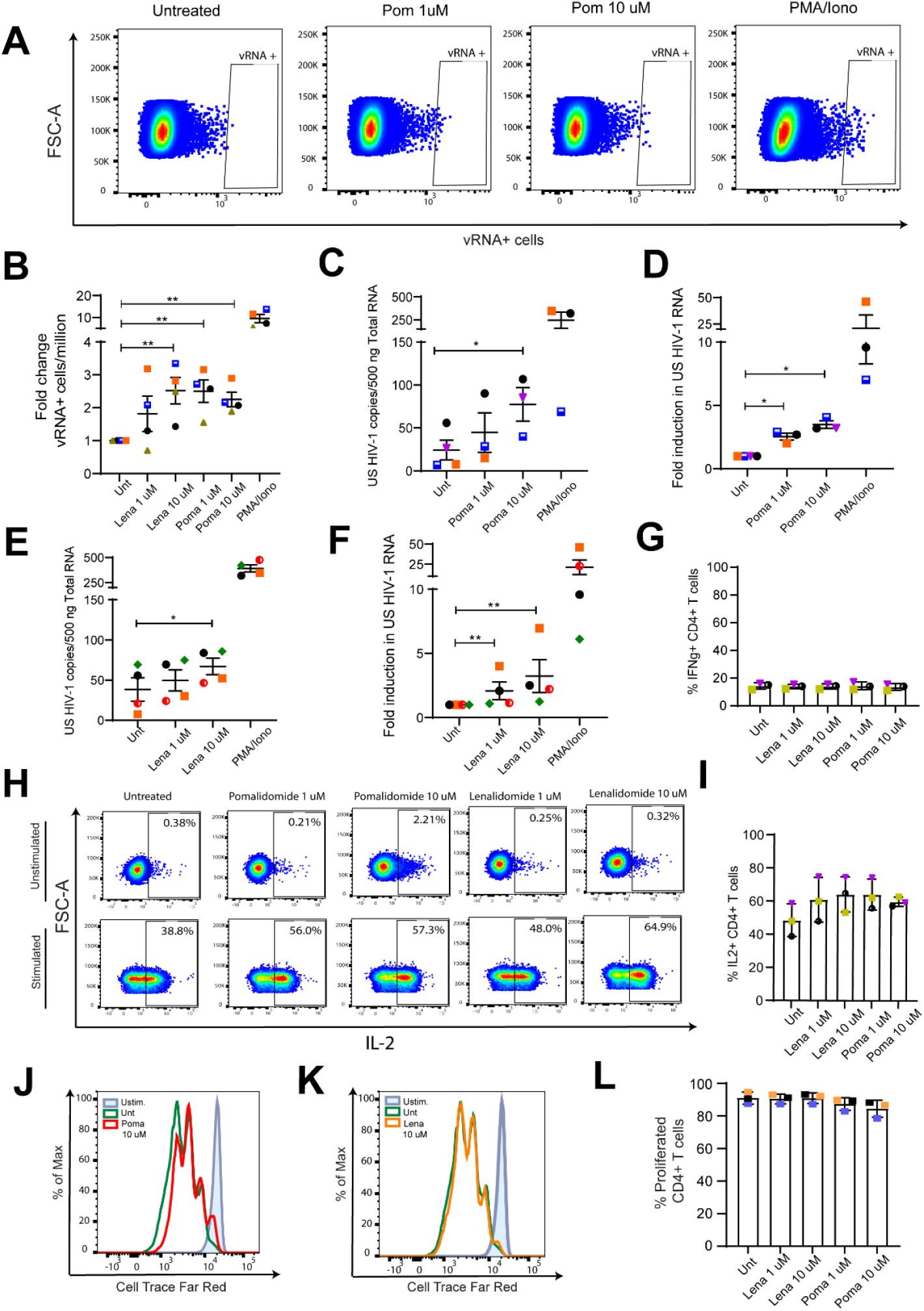
FDA-approved Thalidomide analogues Pomalidomide and Lenalidomide reactivate HIV-1 latency in cells obtained from HIV-1 infected donors. (A) Representative FISH-Flow dot plots of CD4+ T cells from HIV-1 infected donors ex vivo treated with Pomalidomide at 1 µM and 10 µM concentration for 24 hours. (B) Graph panel showing the fold increase in vRNA+ cells/million in ex vivo treated CD4+ T cells from HIV-1 infected donors relative to Untreated control as measured by FISH-Flow (A). Cells were treated with Pomalidomide or Lenalidomide at the indicated concentrations for 24 hours and vRNA+ production was analyzed by FISH-Flow. Error bars represent the mean and standard deviation of independent experiments. Statistical significance was calculated using paired Wilcoxon test, * – p < 0,05, ** – p < 0,01. (C-D) Changes in cell-associated HIV-1 RNA (C) and fold increase (D) in CD4+ T cells isolated from HIV-1 infected donors after treatment with Pomalidomide. CD4+ T cells were isolated from PBMCs from HIV-1 infected donors and treated as indicated for 24 hours. Statistical significance was calculated using paired two-tailed t test (C) and Mann-Whitney test (D), * – p < 0,05, ** – p < 0,01. (E-F) Changes in cell-associated HIV-1 RNA (C) and fold increase (D) in CD4+ T cells isolated from HIV-1 infected donors after treatment with Lenalidomide. CD4+ T cells were isolated from PBMCs from HIV-1 infected donors and treated as indicated for 24 hours. Statistical significance was calculated using paired two-tailed t test (C) and Mann-Whitney test (D), * – p < 0,05, ** – p < 0,01. (G) Representative flow cytometry plots of IFNg production analysis in unstimulated and stimulated primary CD4+ T cells after treatment with Pomalidomide and Lenalidomide. Cells were treated as indicated for 18 hours followed by PMA/Ionomycin (50 ng / 1 µM) stimulation for 7 hours in the presence of a protein transport inhibitor or remained unstimulated. IFN-g production was assessed by intracellular staining and analyzed by flow cytometry. Numbers in the plot show percentage of IFNg producing cells. (H) Percentage of INFg producing CD4+ T cells from 3 healthy donors as described in G (I-J) Representantive histogram of proliferative capacity of unstimulated or aCD3/CD28 stimulated CD4+ T cells in the presence or absence of Pomalidomide (I) or Lenalidomide (J). Cells were stained with a proliferation dye and analyzed 72 hours later by flow cytometry. Dividing cells show decreased intensity of prolifertion dye as it becomes diluted upon cell division. (K) Percentage of proliferated CD4+ T cells from 3 healthy donors in the presence of Pomalidomide and Lenalidomide as described in I-J.

**Table 1.**
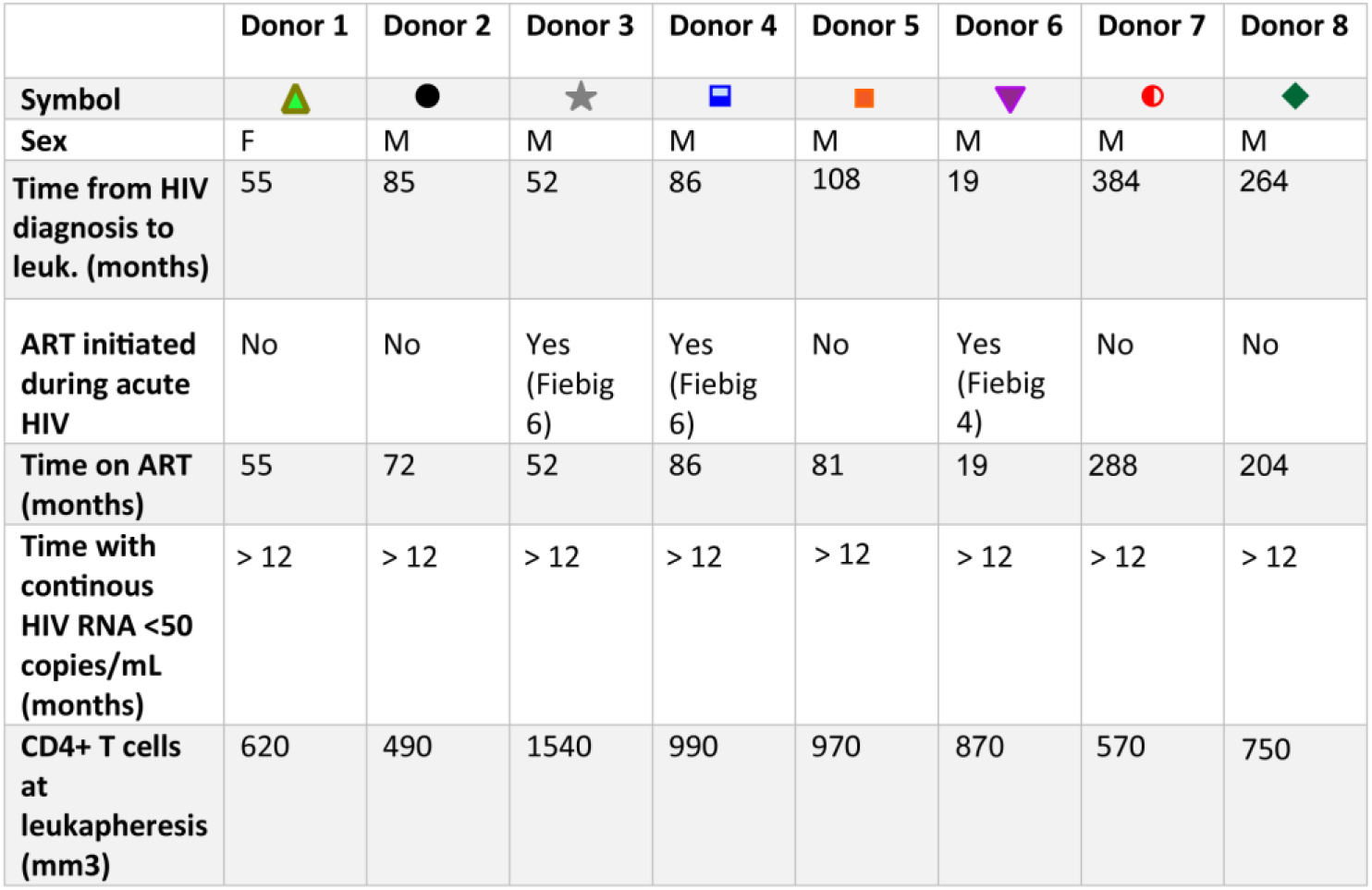
Clinical Information Table corresponding to the HIV-1 infected study participants.

|  | Donor 1 | Donor 2 | Donor 3 | Donor 4 | Donor 5 | Donor 6 | Donor 7 | Donor 8 |
| --- | --- | --- | --- | --- | --- | --- | --- | --- |
| Symbol |  |  |  |  |  |  |  |  |
| Sex | F | M | M | M | M | M | M | M |
| Time from HIV diagnosis to leuk. (months) | 55 | 85 | 52 | 86 | 108 | 19 | 384 | 264 |
| ART initiated during acute HIV | No | No | Yes (Fiebig 6) | Yes (Fiebig 6) | No | Yes (Fiebig 4) | No | No |
| Time on ART (months) | 55 | 72 | 52 | 86 | 81 | 19 | 288 | 204 |
| Time with continous HIV RNA <50 copies/mL (months) | > 12 | > 12 | > 12 | > 12 | > 12 | > 12 | > 12 | > 12 |
| CD4+ T cells at leukapheresis (mm3) | 620 | 490 | 1540 | 990 | 970 | 870 | 570 | 750 |

To assess toxicity induced by treatment with pomalidomide and lenalidomide we treated primary CD4+ T cells obtained from healthy donors with different concentrations of these drugs for 24h and stained for Annexin V as a marker of apoptosis. Besides, we also stained for the extracellular CD69 marker in order to assess global T cell activation caused by thalidomide analogues. Similar to our results with Iberdomide, we show that Pomalidomide and Lenalidomide do not cause loss of viability or apoptosis and do not induce global T cell activation (**Supplementary Figure 12D and E**).

Because of their immunomodulatory role described in literature we also aimed to assess the effects of pomalidomide and lenalidomide treatment in CD4+ T cell proliferation and functionality. As shown with Iberdomide, both Pomalidomide and Lenalidomide treatment did not decrease T cell functionality as measured by intracellular production of IFNg and IL2 upon PMA/Ionomycin stimulation (**Figure 5G-I**), but caused decrease in proliferative capacity of primary CD4+ T cells (**Figure 5J-L**).

Our results show that FDA-approved Pomalidomide and Lenalidomide are able to reverse latency in ex vivo treated CD4+ T cells obtained from PLWH and are therefore promising candidates for potential direct use in the clinic. The immunomodulatory effects of these two drugs together with the fact that they are orally available and have proven to be safe in PLWH, prompts for further studies in the context of HIV-1 latency reversal and cure strategies.

## Discussion

Here, in search of novel determinants of HIV-1 latency, we designed a target discovery strategy based on proteomics, which has allowed us to describe the proteins associated with the HIV-1 5’LTR in its latent and active states. Catchet-MS is based upon a reverse-ChIP strategy and couples CRISPR/dCas9 targeting and purification of a genomic locus to an additional biochemical histone affinity step, to enrich for chromatin-associated dCas9 locus-bound complexes. The use of dCas9 as a bait renders the system suitable for study of single genomic loci in live cells, without the need to modify the site of interest. Additionally, introduction of the histone-based affinity purification step ensures that Catchet-MS effectively removes unwanted background originating from cellular contaminants and non-localized bait molecules from the mass spectrometry analysis. Catchet-MS identified a number of previously identified bona fide interactors of the HIV-1 5’ LTR, as well as a number of novel candidate proteins.

In search of putative novel molecular targets for HIV-1 latency reversal, we focused on factors identified to be significantly enriched or uniquely bound to the latent but not the activated HIV-1 LTR. Using shRNA-mediated depletion, we functionally validated a number of novel putative candidates in J-Lat 11.1 cells. Of 10 putative candidates tested, shRNA depletion of 5 factors (HP1BP3, IKZF1, CDC73, DKC1, PNN) resulted in HIV-1 latency reversal. Of these, the sequence-specific transcription factor IKZF1 stood out as an attractive candidate for which drug targeting is also available (*50, 51*). IKZF1, a critical factor for lymphoid lineage specification in hematopoietic stem cells, suppresses the stem cell and myeloid programs and primes the expression of lymphoid specific genes (*33*). IKZF1 is also an important regulator of T cell function (*40, 60*). Mechanistically, IKZF1 has a dual role in transcription, acting, in a context-dependent manner, both as a repressor and an activator (*32–34*). In an activating capacity, IKZF1 can restrict the activity of the NuRD complex, promoting chromatin accessibility (*34, 36–38*). IKZF1 has also been proposed to act as an adaptor protein for the local recruitment of p-TEFb and the protein phosphatase 1α at IKZF1 target genes, facilitating transcription elongation (*35*). However, in T cells, IKZF1 has been shown to associate with PRC2 and is required for repression of Notch target genes and the hematopoietic stem cell program (*39*). IKZF1-mediated gene silencing may also depend on its interaction with the co-repressors CtBP, CtIP SWI/SNF-related complexes and HDAC-containing Sin3 complexes (*37*).

Consistent with its described repressive mechanisms and functions, depletion of IKZF1 led to a strong reactivation of HIV-1 expression in J-Lat 11.1 and J-Lat A2 cells. Upon PMA-induced transcription, IKZF1 binding downstream of the HIV-1 promoter is abrogated, indicating that IKZF1 binding is required for transcriptional repression. Remarkably, depletion of IKZF1 in J-Lat 11.1 and J-Lat A2 cells reveals that it is required for the establishment of a repressive chromatin environment, characterized by the presence of the repressive chromatin marks H3K9me3 and H3K27me3. Upon IKZF1 depletion the region becomes de-repressed, displaying a drastic loss of enrichment for the H3K27me3 mark, which is even more robust than that observed following PMA-induced LTR activation. Previous evidence points to a model in which, in T-cells, IKZF1 regulates the epigenetic silencing of IKZF1 target genes via recruitment of the PRC2 complex (*39*). The regulatory role of PRC2 at the latent HIV-1 promoter on chromatin compaction, recruitment of PRC1, and repression of HIV-1 transcription is well established (*37, 47, 61*). Consistent with these reports, Catchet-MS identified SUZ12, a core subunit of the PRC2 complex, and CBX8, a subunit of the PRC1 complex, to be associated with the HIV-1 5’LTR in its latent state (Figure 3F). Depletion of IKZF1 led to reduced SUZ12 and CBX8 enrichment over the HS2 region. Our results are consistent with a model in which IKZF1 directly recruits PRC2 to the HIV-1 5’LTR, as previously described at endogenous target genes in T cells (*39*). The PRC2-modified, H3K27 trimethylated region subsequently serves as a docking site for and results in recruitment of PRC1 (**Figure 6**).

**Figure 6.**
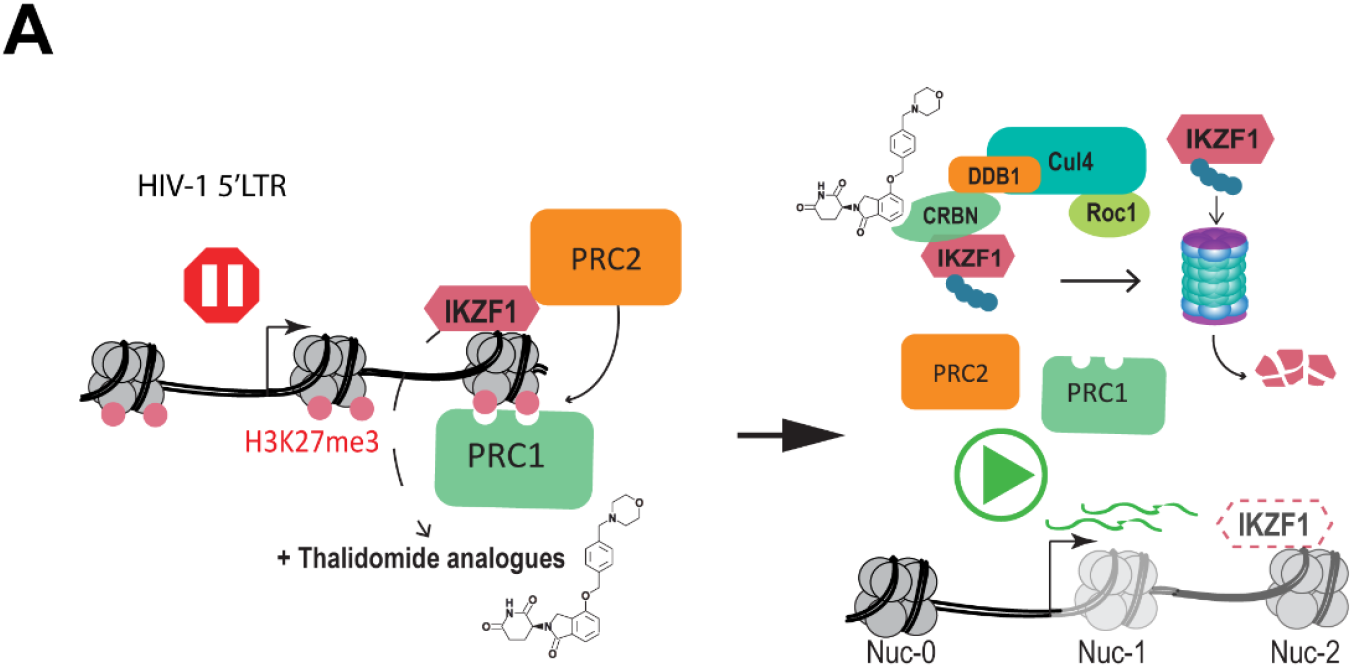
Proposed model for Thalidomide analogues mediated HIV-1 latency reversal. Thalidomide analogues binds to CRBN, a subunit of the CRL^CRBN^ E3 ubiquitin ligase complex that acts as a substrate adaptor. Thalidomide biding induces recruitment of IKZF1 and its ubiquitination by the ligase. In latently infected cells, ubiquitination of IKZF1 and its degradation by the proteasome results in a decrease in IKZF1 levels, impaired LTR recruitment of PRC2 and PRC1, leading to chromatin de-repression and latency reversal.

IMIDs, such as the thalidomide analogues lenalidomide, pomalidomide and iberdomide promote ubiquitin-dependent proteasomal degradation of IKZF1 and IKZF3 by redirecting the substrate specificity of the CRL4^CRBN^ ubiquitin ligase complex (*51, 62*). Among these drugs, iberdomide (CC-220) is a novel compound with the highest reported specificity for CRBN and the lowest IC50, currently in a phase 2 clinical trial for systemic lupus erythematosus (SLE) and multiple myeloma (MM). Due to its high potency we selected iberdomide for treatment of CD4+ T cells obtained from cART suppressed aviremic participants. Iberdomide mediated depletion of IKZF1 was accompanied by pronounced reversal of HIV-1 latency, with no significant associated toxicity. BET inhibitors, such as JQ1, represent a class of compounds previously reported to synergize with IMIDs in inhibiting the growth of refractory forms of MM (*53, 54*). As BET inhibitors are also a well-established class of HIV-1 LRAs, we examined the latency reversal effects of combination treatment with iberdomide and JQ1. Remarkably, co-treatment of latently infected primary CD4+ T cells results in strong and synergistic induction of HIV-1 transcription in cells obtained from HIV infected cART suppressed aviremic patients. Thus, similar to what was observed in the treatment of resistant MM, combination of these two classes of compounds also leads to synergism in the context of HIV-1 latency reversal.

Although clinically advanced, Iberdomide is yet to be approved by international drug approval bodies. On the other hand, Pomalidomide and Lenalidomide are FDA-approved compounds currently in use for the treatment of immune disorders such as MM or other lymphopathies (*57*); therefore, we aimed to investigate their role in HIV-1 latency reversal and potential effects on the immune system. We show that both Pomalidomide and Lenalidomide treatment of CD4+ T cells obtained from HIV-1 infected, aviremic donors leads to reactivation of HIV-1 latency. Importantly, we show that high concentrations of Pomalidomide and Lenalidomide do not cause undesirable associated toxicity or global immune activation in primary CD4+ T cells.

The role of Thalidomide and its analogues in modulating the immune system has been very well studied and described in the past, in the context of lympho and myeloproliferative immune pathologies (*57*). Consistent with what is reported in the literature (*57*), in this study we show that Iberdomide and the FDA-approved Thalidomide analogue Pomalidomide modulate the proliferation capacity of primary T cells. In addition, we demonstrate that the use of Iberdomide does not affect T cell functionality, as measured by cytokine expression in CD4+ and CD8+ T cells and does not cause apoptosis or global immune activation. More importantly, high concentrations of Pomalidomide and Lenalidomide do not cause cytotoxicity or impair T cell functional response in primary CD4+ T cells obtained from healthy donors. Crucial to the potential applicability of LRA treatments alone or in combination in future clinical studies is the absence of detrimental effects on the overall T cell biology and the preservation of a functional cytotoxic compartment. Currently available LRAs, though not appearing to cause serious adverse effects in HIV-1 infected individuals, lead to significant cytotoxicity in CD4+ and CD8+ T cells and thus major impairment of cytotoxic immune responses (*22*). As viral clearance after latency reactivation depends mostly on cytotoxic CD8+ T cells, maintaining or boosting functional immune responses in HIV-1 cure strategies is of utmost importance for achieving long-term reduction in reservoir size. Thus, the role of Thalidomide analogues in modulating the immune system together with their LRA activity makes them of great interest for HIV-1 cure studies. In fact, previous clinical studies have shown that the use of Pomalidomide leads to an increase in the total numbers of CD4+ and CD8+ T cells, decrease in senescence and a shift towards central memory T cells in HIV-1 infected individuals. Furthermore, IMIDs such as Pomalidomide have been shown to enhance co-stimulation of virus specific CD8+ T cells (*63*) and, more importantly, treatment of Pomalidomide and Lenalidomide significantly expands and enhances the effector function of HIV-1 specific CD8+ T cells in vitro (*64*) and boosts immune responses in HIV-1 infected individuals by inducing chemotaxis-mediated recruitment of effector cells (*65*). Notably, several clinical trials have shown that Pomalidomide and Lenalidomide at high doses are safe and tolerable in HIV-1 infected patients in the context of AIDS-related morbidities, such as Kaposi sarcoma, or in HIV+ individuals suffering from lymphoma (*59, 66–68*).

To conclude, FDA/EMA-approved Thalidomide analogue compounds, such as Pomalidomide and Lenalidomide, which target IKZF1 for degradation and lead to HIV-1 latency reactivation while boosting HIV-1 specific CTL responses, represent attractive candidates for inclusion in proof of concept clinical studies aiming to reduce the latent HIV-1 reservoir. The specificity of the IMID/IKZF1-targeting pharmacological strategy for HIV-1 latency reversal may also be enhanced by combination treatment with other LRA class compounds, such as BET inhibitors, which we demonstrate to lead to synergistic latency reversal.

## Materials and Methods

### Cell lines

Latent HIV-1 infected Jurkat cells (clone J-Lat 11.1 and clone J-Lat A2) were cultured in RPMI-1640 media supplemented with 7% FBS and 100 µg/ml penicillin-streptomycin at 37°C in a humidified 95% air, 5% CO2 incubator. Primary CD4+ T cells were cultured in RPMI-1640 media supplemented with 7% FBS and 100 µg/ml penicillin-streptomycin at 37°C in a humidified 95%-air-5%CO2 incubator. HEK 293T cells were cultured in DMEM supplemented with 10% FBS and 100 µg/ml penicillin-streptomycin at 37 °C in a humidified 95% air 5% CO2 incubator and used exclusively for the production of lentiviral shRNA vectors.

### Establishment of J-lat 11.1 cells expressing the HA-V5-FLAG-dCas9 bait

In order to generate J-Lat 11.1 cells expressing the triple tagged (HA, V5, 3xFLAG) dCas9 construct, cells were then nucleofected with a modified version of the PX462 plasmid (AddGene) and selected for resistance to puromycin. pSpCas9n(BB)-2A-Puro (PX462) V2.0 was a gift from Feng Zhang (Addgene plasmid # 62987 ; http://n2t.net/addgene:62987 ; RRID:Addgene_62987). The original plasmid encodes for a Cas9 D10A nickase mutant, we modified the original sequence by introducing an H841A mutation that renders the Cas9 fully catalytically inactive (dCas9). Additionally, we introduced a triple FLAG epitope sequence (3xFLAG), a V5 epitope sequence and an HA epitope sequence at the N-terminal region of the dCas9. The plasmid contains a chicken β-actin promoter that drives the expression of the dCas9 bait and a U6 promoter that drives the expression of the gRNA scaffold. The sequence of the gRNA is 5’-GAAGCGCGCACGGCAAG-3’ and is located between nucleotides 708 and 724 on the HIV-1 5’ LTR, downstream the transcriptional start site (TSS) and in-between Nuc-1 and Nuc-2 (Figure 1A). Within the plasmid, the dCas9 open reading frame (ORF) sequence additionally contains an IRES sequence that is followed by a puromycin resistance cassette, allowing for puromycin selection of dCas9 expressing cells. The plasmid sequence information is available upon request. 48 hours after nucleofection the cells were selected for a week with 1μg/ml puromycin and expanded in a polyclonal population. Unstimulated J-Lat 11.1 cells were used to model the latent, repressed, promoter while cells stimulated with 10 μm of phorbol 12 myristate 13-acetate (PMA) (Sigma), a potent PKC agonist, were used to model the transcriptionally active promoter.

### Plasmids nucleofection

The expression plasmids for the HA-V5-FLAG-dCas9 bait and the dCas9-VPR were delivered to J-Lat 11.1 cells by nucleofection using Amaxa Nucleofector (Lonza) as previously described (*3*) and the Cell Line Nucleofector Kit R (Lonza). Briefly, cells were split to 4 × 10^5^ cells/ml one day before nucleofection, 8 million cells were centrifuged at 600rcf for 5 min at room temperature, resuspended in 100 μl of solution R, and nucleofected with 2 μgs of plasmid, using program O28 and the Cell line nucleofector Kit R. Nucleofected cells were then resuspended in 500 μl of pre-warmed, serum-free antibiotic-free RPMI at 37 °C for 15 min and then plated in 5 ml of pre-warmed complete media. Seventy-two hours post-nucleofection cells nucleofected with the dCas9 VPR plasmid were analyzed with flow cytometry while cells nucleofected with the dCas9 construct for purification (HA, V5, FLAG) were selected for 10 days in puromycin 0.5μg/ml and expanded as a polyclonal population.

### Bacterial strains

DH5α and Stable3 bacterial cells were used to propagate the plasmids used in the study.

### Western Blotting

10x10^6^ cells were lysed with 100 µl cold lysis buffer containing 150 mM NaCl, 30 mM Tris (pH 7.5), 1 mM EDTA, 1% Triton X-100, 10% Glycerol, 0.5 mM DTT, protease inhibitor cocktail tablets (EDTA-free) (Roche) at 4°C per 30 minutes. Cell lysates were clarified by centrifugation (14,000 rpm for 30 min at 4°C), mixed with 4x sodium dodecyl sulfate (SDS)-loading buffer with 0.1M DTT and boiled at 95°C per 5 min. Samples were run in a 10% SDS-polyacrylamide gel at 100V. The proteins were then transferred to polyvinylidene difluoride (PVDF) membranes (Immobilion) and the membranes probed with primary antibodies (anti-V5 epitope (Invitrogen), anti-HA epitope (Abcam), anti-FLAG epitope (Sigma) and anti-tubulin (Abcam) overnight at 4 °C. Secondary antibodies were used accordingly to the species reactivity and incubated for 1 hour at room temperature. Proteins were detected by chemical luminescence using the SuperSignal West Pico or SuperSignal West Pico Femto Chemiluminescent Substrate (Thermo Scientific). For visualization of dCas9 protein enrichment during the ChIP-MS protocol, the protein complexes bound to the beads were decrosslinked and eluted and at 95°C for 30min in 4x sodium dodecyl sulfate (SDS)-loading buffer containing 0.1M DTT. The proteins were then resolved in a 10% polyacrylamide SDS-PAGE gel and western blotting was performed using a mouse anti-FLAG antibody, a secondary anti-mouse HRP antibody, and imaged as above.

### Cytoplasmic and nuclear fractionation

Approximately 2x10^7^ J-lat 11.1 cells expressing the HA-V5-FLAG-dCas9 bait were subjected to cellular fractionation to separate the cytoplasmic and nuclear fraction. We adapted the cellular fractionation protocol described in (Ten Have et al., 2012) to cells in suspension. Briefly, the cells were collected by centrifugation at 800rcf for 5 minutes at 4C and washed twice with PBS. The cell pellet was then transferred and resuspended into a pre-chilled 7ml dounce homogenizer, where the cell membrane was broken using 10 strokes of a tight pestle. The dounced cells were then centrifuged for 5 minutes at 4C, 800rcf and the supernatant was retained as cytoplasmic fraction. The nuclear pellets were then resuspended in 3ml of a solution (S1) containing 0.25M Sucrose and 10mM MgCl2 and layered over a 3ml cushion of a 0.35M Sucrose and 0.5mM MgCl2 solution (S2) by slowly pipetting S1 on top S2. The cell pellets were then centrifuged again for 10 minutes at 4C at 2500rcf and the remaining pellets retained as the nuclear fraction.

### Chromatin immunoprecipitation and HIV-1 5’LTR purification using Catchet-MS

In order to more specifically enrich for chromatin and reduce background from cellular contaminants and from the non-chromatin associated bait, we improved the stringency of our in-house ChIP protocol (*3*) by incorporating a few steps from the Chromatin enrichment for proteomics (CheP) (*31*) during the preparation of our input material. We have then followed a traditional ChIP protocol in which we have enriched for the dCas9 bait and its associated complexes using an Anti-V5 Agarose Affinity Gel (Sigma). To further eliminate the background originating from the purification of non-chromatin associated bait complexes we have additionally added a histone enrichment step, downstream of the bait enrichment step, using histone H2B (Abcam) and H3 antibodies (Abcam) conjugated to Protein G SepharoseTM 4 Fast flow (GE Healthcare) and Protein A SepharoseTM 4 Fast flow (GE Healthcare) beads by dimethyl pimelimidate cross-linking as described in (*69*).

### Chromatin input preparation using CheIP

Approximately 3 billion cells per condition were collected, cross-linked and processed in batches of 400million cells. After collection, by centrifugation at 800rcf for 5 minutes, each batch was washed two times in 40 ml PBS supplemented with 1mM MgCl2 and 1mM CaCl2, and cross-linked in 40 ml PBS with 1% formaldehyde (Formaldehyde methanol free 16%, Polysciences inc.) at room temperature for 30 min with vertical rotation at 15rpm. The reaction was then quenched with the addition of 1 M glycine to a final concentration of 125 mM. Cross-linked cells pellet were then washed in 20 ml cold buffer B (0.25% Triton x-100, 1mM EDTA, 0.5mM EGTA, 20mM HEPES pH 7.6) and 20 ml cold buffer C (0.15M NaCl, 1mM EDTA, 0.5mM EGTA, 20mM HEPES pH 7.6), resuspended in 2 ml ChIP incubation buffer (1% Triton x-100, 150mM NaCl, 1mM EDTA, ph 8.0, 0.5 mM EGTA, 20mM HEPES pH 7.6, protease inhibitor cocktail tablets (EDTA-free, Roche) and then transferred in 15ml polystyrene Falcon tubes compatible with sonication. In order to isolate and expose the cross-linked nuclei to the subsequent denaturation steps the samples were sonicated with 15 ml probes at medium intensity for 10 minutes (30’ ON/30’ OFF intervals; Diogenode Bioruptor) and the cell nuclei were spun at 800rcf for 10 minutes. The pellets were then re-suspended in 2 ml ChIP incubation buffer containing 0.15% SDS and sonicated again for 10 minutes at medium intensity (30’ ON/30’ OFF intervals; Diogenode Bioruptor). The sonicated nuclei in suspension were then aliquoted (500µls per tube) in 2ml Eppendorf tubes and denatured for 5 minutes at room temperature with 350µls 10% SDS and 1.2 ml 8M Urea buffer (10mM Tris pH 7.4, 1mM EDTA and 8 M urea). The insoluble fraction of the nuclei, enriched for chromatin, was then precipitated by centrifugation at 16200rcf at room temperature for 20minutes. The precipitated pellets were then resuspended and combined (4 into 1) in a single 2ml Falcon tube and denatured again for 5 minutes at room temperature with 500µls of 4% SDS buffer (50mM Tris ph 7.4, 10 mM EDTA, 4% SDS) and 8M Urea buffer and then precipitated again by centrifugation at 16200rcf at room temperature for 20minutes.

The pellets were then resuspended and washed by centrifugation at 16200rcf at room temperature for 20minutes in 2ml 4% SDS buffer and then resuspended in 2.5 ml ChIP incubation buffer containing 1% SDS. The content of each tube was then transferred in a 15ml polystyrene tube and sonicated (3 tubes at the time) at a high intensity (30’ ON/30’ OFF intervals; Diogenode Bioruptor), using metal probes, to obtain chromatin fragments in a 200-500bp size range. The different chromatin batches were then pulled together concentrated through 100kDa Centricon filters, to buffer exchange with 0.1% SDS ChIP incubation buffer and remove smaller proteins and fragments. The sonicated chromatin (a total of 50 ml per condition) was then spun at max speed (20817 rcf) at 4°C for 30 minutes to remove cellular debris and diluted to 50 ml per condition. The chromatin was then precleared overnight at 4°C with vertical rotation at 15rpm (a total of 5 reactions per condition in 15ml tubes) with 500µls of a mix of Protein G and A Sepharose^TM^ 4 Fast flow (GE Healthcare) beads and then used for immunoprecipitation.

### HA-V5-FLAG-dCas9 bait enrichment

Per each condition, dCas9 containing complexes were isolated from the sonicated chromatin using 500µls (a total of 5 reaction where 100µls of affinity beads were used to purify 10ml of chromatin) of Anti-V5 Agarose Affinity Gel (Sigma) and then washed two times per each buffer (5minutes, 15rpm vertical rotation washes) with buffer 1 (0.1% SDS, 0.1% DOC, 1% Triton x-100, 150mM NaCl, 1 mM EDTA pH 8.0, 0.5 mM EGTA, 20mM HEPES pH8.0.), buffer 2 (500mM NaCl: 0.1% SDS, 0.1% DOC, 1% Triton x-100, 500mM NaCl, 1 mM EDTA pH 8.0, 0.5mM EGTA, 20mM HEPES pH8.0), buffer 3 (0.25M LiCL, 0.5%DOC, 0.5% NP-40, 1mM EDTA pH 8.0, 0.5mM EGTA pH 8.0, 20mM HEPES pH 8.0), buffer 4 (1mM EDTA pH8.0, 0.5mM EGTA pH 8.0, 20mM HEPES pH 8.0) to remove unspecific binding and eluted in a total volume of 1 ml elution buffer 1% SDS, 0.1M NaHCO3 per each condition. At this stage a 50µl aliquot of the eluate was taken to be analyzed by qPCR as quality control for dCas9 enrichment over the chromatin input at the desired chromatin locus.

### Locus enrichment

The eluate was then diluted 10 times in ChIP incubation buffer without SDS (1% Triton x-100, 150mM NaCl, 1mM EDTA, ph 8.0, 0.5 mM EGTA, 20mM HEPES pH 7.6, protease inhibitor cocktail tablets (EDTA-free) (Roche) and immunoprecipitated again in a single reaction with 100µls of a mix of Protein G and A SepharoseTM 4 Fast flow (GE Healthcare) pre-conjugated with histone H3 and histone H2B antibodies in order to specifically enrich for chromatin-associated dCas9 complexes and eliminate from the purification unspecific non-chromatin associated dCas9 complexes. The immunoprecipitated material, bound to the beads, was then washed again 2 times per each buffer with buffer 1, buffer 2, buffer 3 and buffer 4 to remove unspecific binding and the protein complexes bound to the beads were finally eluted and decrosslinked by boiling the sample for 30min in 100µls of 4x sodium dodecyl sulfate (SDS)-loading buffer, containing 0.1M DTT.

### Mass spectrometry

Proteins were resolved in a 15% polyacrylamide SDS-PAGE gel, visualized by Colloidal Blue Staining Kit (ThermoFisher) and prepared for nanoflow LC-MS/MS analysis. SDS-PAGE gel lanes were cut into 1-mm slices and protein were subjected to in-gel reduction with dithiothreitol, alkylation with iodoacetamide and digested with trypsin (TPCK Trypsin, ThermoScientific, Rockford, IL, USA), as described previously (*70*). Peptides were extracted from the gel blocks in 30 % acetonitrile (Biosolve) and 0.5 % formic acid (Biosolve) and dried in a SpeedVac Vacuum Concentrator (ThermoFisher). Samples were analyzed by LC-MS/MS on a 20 cm x 75 µm C18 column (BEH C18, 130 Å, 3.5 µm, Waters) after trapping on a nanoAcquity UPLC Symmetry C18 trapping column (Waters, 100 Å, 5 µm, 180 µm x 20 mm) on an EASY-nLC 1000 coupled to a Fusion Tribrid Orbitrap mass spectrometer (ThermoFisher Scientific), essentially as described in (*71*). Data analysis was performed with the Mascot software suite (Daemon version 2.3.2 and Distiller version 2.3.02, Matrix Science) within the Proteome Discoverer (version 2.2, ThermoFisher Scientific) framework. Spectra were searched against a Uniprot database (version April 2017, taxonomy Homo sapiens) concatenated with a fasta database containing amino acid sequences of the triple tagged dCas9 (HA, V5, 3xFLAG) construct. Protein Mascot scores and peptide numbers were taken directly from the Mascot output and reported. The same procedure was used to analyze input samples for the protocol characterization (Supplementary Figure 3). Heat maps were generated using MORPHEUS (http://software.broadintistute.org/morpheus/index.html).

### Chromatin quality control

100µl aliquots of chromatin were taken per each batch of the purification protocol, resuspended in 500µls of ChIP elution buffer and, after addition of 24µl 5M NaCl (200mM final concentration), decrosslinked overnight at 65°C on a heat block. DNA isolation was then performed using phenol-chloroform isoamyl isolation method and ethanol precipitation in the presence of glycogen as a carrier. The DNA pellet was resuspended in 100µls of nuclease-free water. 5µls of DNA was then mixed with sample 4x sample buffer and run on a 1% agarose DNA electrophoreses gel to check for fragmentation between 200 and 500bps.

### ChIP-qPCR

Real-time qPCR was used to detect the specific enrichment of the 5’-LTR locus at a DNA level after the ChIP procedure. Reactions of 10 µl were performed with the GoTaq qPCR Master Mix kit (Promega) in a CFX Connect Real-Time PCR thermocycler (Biorad) Relative quantitation was calculated with the 2-ΔCt method where relative enrichment over the adjusted DNA input was used for data representation.

To evaluate IKZF1, CBX8 and SUZ12 binding to the HIV-1 promoter and histone marks profile a smaller scale ChIP experiment (50 million cells) with identical preparation of the chromatin input as for the mass spectrometry experiment. For the immunoprecipitation, protein G-beads or A-beads were used in combination with the anti-IKZF1 antibody (Cell signal technology), anti-SUZ12 (abcam), anti CBX8 (homemade) and histone antibodies: Histone H3 (abcam), histone H3K27me3 (Millipore), H3K4me3 (Millipore), H3K9me3 (Abcam). DNA was PCI extracted and used to evaluate the enrichment of ChIP-Ikaros at the HIV-1 5’LTR by the same primer sets summarized in the key resources table. Antibodies used are summarized in the Key resource table.

### ChIP-Sequencing data analysis

In order to align the short reads produced by the ChIP-Seq procedure and also capture reads mapped to the HIV genome, we constructed a custom version of the human genome (UCSC version hg38) where we attached the HIV genome as an additional chromosome. We used the HIV strain K03455.1 (https://www.ncbi.nlm.nih.gov/nuccore/K03455). Subsequently, the resulting genome was indexed and reads were aligned with bwa 0.7.17 (1) and the bwa mem algorithm. In order to compensate for differences in library sizes between ChIP and Input DNA sequencing samples, the total number of reads of each sample was equalized by uniformly downsampling reads relatively to the sample with the lower number of reads. ChIP peaks were then called with MACS2 (2) using default parameters. The peaks returned by MACS2 were further filtered by i) imposing an additional MACS FDR threshold of 0.1%, ii) excluding peaks demonstrating a fold enrichment less than 3 in log2 scale (where fold enrichment is the ratio of normalized reads under a peak area in the ChIP sample to the respective number of normalized reads in the Input DNA sample). Finally, in order to visualize the ChIP-Seq signals, we constructed a custom UCSC track hub based on the combined hg38-HIV genomes using instructions and tools provided by UCSC (*72, 73*).

### Production of shRNA lentiviral vectors

Lentiviral constructs containing the desired shRNA sequence were amplified from bacterial glycerol stocks obtained in house from the Erasmus Center for Biomics and part of the MISSION® shRNA library (sigma). Briefly, 5.0 x 10^6^ HEK293T cells were plated in a 10 cm dish and transfected with 12,5 μg of plasmids mix. 4,5 μg of pCMVΔR8.9 (envelope), 2 μg of pCMV-VSV-G (packaging) and 6ug of shRNA vector were mixed in 500 μL serum-free DMEM and combined with 500 μL DMEM containing 125 μL of 10 mM polyethyleneimine (PEI, Sigma). The resulting 1 mL mixture was applied to HEK293T cells after 15 min incubation at room temperature. The transfection mix was removed after 12 hr and replaced with fresh RPMI medium. Virus-containing medium was harvested twice and replaced with fresh medium at 36 hours 48 hr, 60, 72 hr post-transfection. After each harvest, the collected medium was filtered through a cellulose acetate membrane (0.45 μm pore) and used directly for shRNA infections or stored at -80°C for subsequent use. Lentivaral vectors used in the study are summarized in the summarized the key resources table.

### Flow Cytometry of J-lat 11.1

GFP expression in the J-Lat cell lines was analyzed by Flow Cytometry. The live-cell population was defined by using the forward scatter area versus side scatter area profiles (FSC-A, SSC-A). Single cells were then gated by using forward scatter height (FSC-H) versus forward scatter width (FSC-W) and side scatter height (SSC-H) versus side scatter width (SSC-W). GFP intensity to differentiate between GFP-positive and negative cells. Between 2 - 4x105 events were collected per sample on an LSR Fortessa (BD Biosciences) and analyzed using FlowJo software (version 9.7.4, Tree Star).

### Fluorescence microscopy

Fluorescence microscopy and bright field pictures of J-Lat 11.1 cells in the absence and presence of PMA were acquired using an Olympus IX70 Fluorescence Phase Contrast inverted Microscope.

### Total RNA Isolation and quantitative real-time PCR (RT-qPCR)

Total RNA was isolated from the cells using Trizol (Thermo Fisher) and the Total RNA Zol out kit (A&A Biotechnology) and residual genomic DNA digested with DNAseI (Life technologies). cDNA synthesis reactions were performed with Superscript II Reverse Transcriptase (Life Technologies) kit following with random priming and manufacturer’s instructions. RT-qPCR reactions were conducted CFX Connect Real-Time PCR Detection System thermocycler (BioRad) using GoTaq qPCR Master Mix (Promega) following manufacturer protocol. Amplification was performed on the using following thermal program starting with 3 min at 95 °C, followed by 40 cycles of 95 °C for 10 s and 60 °C for 30 s. The specificity of the RT-qPCR products was assessed by melting curve analysis. Primers used for real-time PCR are listed in Table 1. Expression data was calculated using 2-ΔΔCt method by Livak Schmittgen (*74*). Cyclophilin A (CyPA), GAPDH and β2 microglobulin (B2M) were used as housekeeping genes for the analysis.

### Isolation and ex vivo infection of primary CD4+ T cells

Peripheral blood mononuclear cells (PBMCs) were isolated by Ficoll density gradient sedimentation of buffy coats from healthy donors (GE Healthcare). Total CD4+ T cells were separated by negative selection using EasySep Human CD4+ T Cell Enrichment Cocktail (StemCell Technologies). Primary CD4+ T cells were plated at a concentration of 1,5x10 ^6^ cells/mL left overnight for recovery. HIV-1 latency *ex vivo* model was generated by spinoculation according to Lassen and Greene method as described elsewhere (*52*). Pseudotyped viral particles were obtained from co-transfecting HEK 293T cells with HXB2 Env and pNL4.3.Luc.R-E-plasmids using PEI. Supernatants were collected at 48 and 72h post-transfection, filtered with a 0.45 µM filter and stored at -80°C. Expression vectors HXB2 Env and pNL4.3.Luc.R-E-were donated by Dr. Nathaniel Landau and Drs. Kathleen Page and Dan Littman, respectively. Antiretroviral drugs Saquinavir Mesylate and Raltegravir were kindly provided by the Centre for AIDS reagents, NIBSC.

### Flow cytometry for T cells activation and toxicity assay

Primary CD4+ T cells isolated from buffy coats of healthy volunteers were treated with 10uM of iberdomide, 500nM of JQ1, or both compounds, with PMA/ionomycin used as a positive control. The cells were examined by flow-cytometry at 24 and 48 hours, live cells were gated using forward scatter area versus side scatter area profiles (FSC-A, SSC-A). Cells were then stained for expression of Annexin V to examine the percentage of cells undergoing apoptosis and with the surface receptor CD69 to measure T cells activation. For Annexin V and CD69 staining, 10^6^ cells were washed with PBS supplemented with 3% FBS and 2.5 mM CaCl2 followed by staining with Annexin V-PE (Becton and Dickinson), CD69-FITC (eBIOSCIENCE) for 20 minutes at 4C in the presence of 2.5 mM CaCl2. Cells were then washed with PBS/FBS/CaCl2 and analyzed by flow cytometry. Between 2 - 4x10^5^ events were collected per sample within 3 hours after staining on an LSRFortessa (BD Biosciences, 4 lasers, 18 parameters) and analyzed using FlowJo software (version 9.7.4, Tree Star).

### T cell proliferation and functionality assay

To analyze the effect of the LRA on CD8+ and CD4+ T cells, proliferation and cytokine expression were analyzed by flow cytometry. Primary CD8+ and CD4+ T cells were isolated from buffy coats form healthy donors by Ficoll gradient (GE Healthcare) followed by negative selection with RosetteSep Human CD8+ T Cell Enrichment Cocktail or RosetteSep Human CD4+ T Cell Enrichment Cocktail (StemCell Technologies) respectively. Isolated cells were left overnight for recovery. To analyze T cell proliferation capacity, 1 million CD8+ or CD4+ T cells were stained with 0,07 uM CellTrace Far Red Cell Proliferation dye (ThermoFisher Scientific) following manufacturer’s instructions. Cells were then cultivated for 72 hours with either unstimulated or stimulated conditions in the presence of the LRA, and analyzed by flow cytometry. Stimulation of T cells was performed using Anti-CD3/CD28 coated Dynabeads (ThermoFisher Scientific) following manufacturer’s protocol. Proliferation capacity was determined by a decrease in proliferation dye intensity in daughter cells upon cell division. To analyze T cell functionality by means of cytokine expression 1 million CD8+ or CD4+ T cells were left untreated or treated with the LRA for 18 hours. Cells were then left unstimulated or stimulated with 50 ng/mL PMA and 1uM Ionomycin for 7 hours in the presence of a protein transport inhibitor (BD GolgiPlug^TM^, BD Biosciences). To stain for intracellular cytokines cells were washed with PBS supplemented with 3% FBS followed by a fixation and permeabilization step with FIX & PERM Kit (Invitrogen) following manufacturer’s protocol and incubated with 1:25 BV510 anti-IL2 (563265, BD Biosciences) and PE-Cy7 anti-IFNg (27-7319-41, eBioscience) in permeabilization buffer for 45 minutes at 4C. Stained cells were washed with PBS supplemented with 3% FBS and analyzed by flow cytometry.

### HIV-1 latency reversal in primary CD4+ T cells isolated from aviremic patients

Primary CD4+ T cells from 5 aviremic patients (maintained viremia below 50 copies/mL for at least two years) were isolated as described previously (*75*). Two million CD4+ T cells were plated at the cell density of 10^6^/ml and treated as indicated. After 16 hours cells were lysed in TRI reagent and RNA was isolated with Total RNA Zol-out kit (A&A Biotechnology), cDNA synthesis was performed as described above. Absolute quantification of cell-associated HIV-1 gag RNA was performed in a nested PCR as described previously (*76*). Briefly, first round of amplification was performed in a final volume of 25 μl using 4 μl of cDNA, 2.5 μl of 10× PCR buffer (Life Technologies), 1 μl of 50mM MgCl2 (Life Technologies), 1 μl of 10 mM dNTPs (Life Technologies), 0.075 μl of 100 μM Gag Forward primer, 0.075 of 100 μM SK437 Reverse primer, and 0.2 μl Platinum Taq polymerase (Life Technologies). The second round of amplification was performed in a final volume of 25 μl using 2 μl of pre-amplified cDNA, 2.5 μl of 10× PCR buffer (Life Technologies), 1 μl of 50mM MgCl2 (Life Technologies), 1 μl of 10 mM dNTPs (Life Technologies), 0.05 μl of 100 μM Gag Forward primer, 0.05 μl of 100 μM Gag Reverse primer, 0.075 μl of 50 μM Gag Probe and 0.2 μl Platinum Taq polymerase. The absolute number of gag copies in the PCR was calculated using a standard curve ranging from 8 to 512 copies of a plasmid containing the full-length HIV-1 genome (pNL4.3.Luc.R-E-). The amount of HIV-1 cellular associated RNA was expressed as number of copies/μg of input RNA in reverse transcription. The study was conducted according to the ethical principles of the Declaration of Helsinki. All patients involved in the study gave their written informed consent. The study protocol and any amendment were approved by The Netherlands Medical Ethics Committee (MEC-2012-583).

### FISH-Flow analysis

Cells were collected, fixed, permeabilised and subjected to the PrimeFlow RNA assay (Thermo Fisher Scientific) following the manufacturer’s instructions and as described in (Rao, 2021, Nat Comm). Primary CD4+ T cells were first stained in Fixable Viability dye 780 (Thermo Fisher Scientific) for 20 minutes at room temperature (1:1000 in dPBS). HIV unspliced mRNA was thenlabelled with a set of 40 probe pairs against the GagPol region of the vRNA (catalogue number GagPol HIV-1 VF10-10884, Thermo Fisher Scientific) diluted 1:5 in diluent provided in the kit and hybridized to the target mRNA for 2 hours at 40°C. Samples were washed to remove excess probes and stored overnight in the presence of RNAsin. Signal amplification was then performed by sequential 1.5 hours, 40°C incubations with the pre-amplification and amplification mix. Amplified mRNA was labelled with fluorescently-tagged probes for 1 hour at 40°C. Samples were acquired on a BD LSR Fortessa Analyzer and gates were set using the uninfected CD4+ T cells. The analysis was performed using the FlowJo V10 software (Treestar).

### Statistical analysis

Statistical analysis was conducted as indicated in the figure legends using Prism version 8.3.0 (GraphPad software).

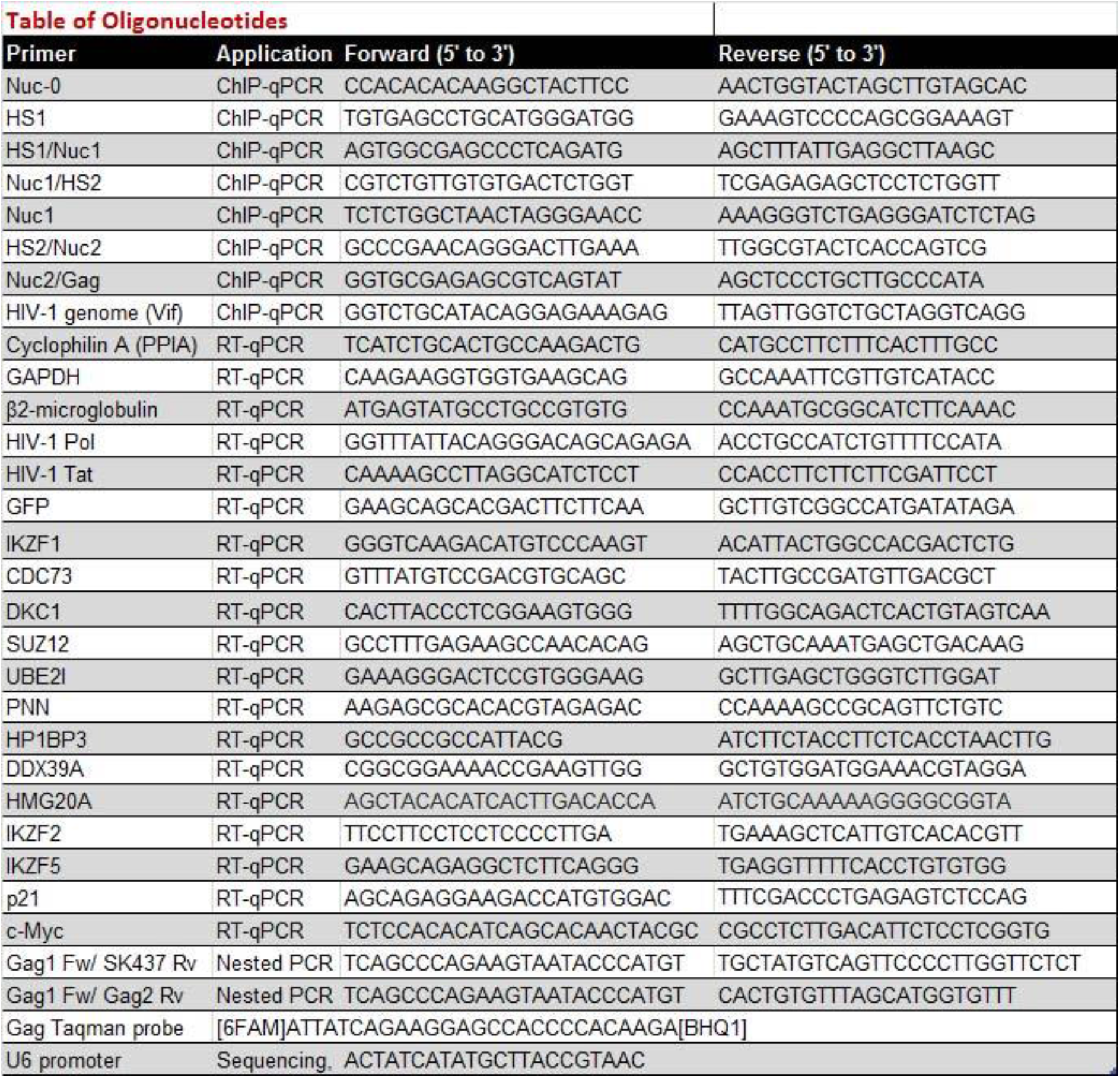

## Acknowledgments

We thank Vaggelis Harokopos of the BSRC Al. Fleming Genomics Facility for NGS.

## Author Contributions

EN, RC, SR, RI, SK, TS, DD, AG, and RJP carried out experiments and performed data analysis. PM, and PH performed Nextgen sequencing and data analysis. EN, RC, PH, RJP, JD and TM conceived the study and wrote the manuscript. All authors read and approved the final manuscript.

---

**Figure S1.**
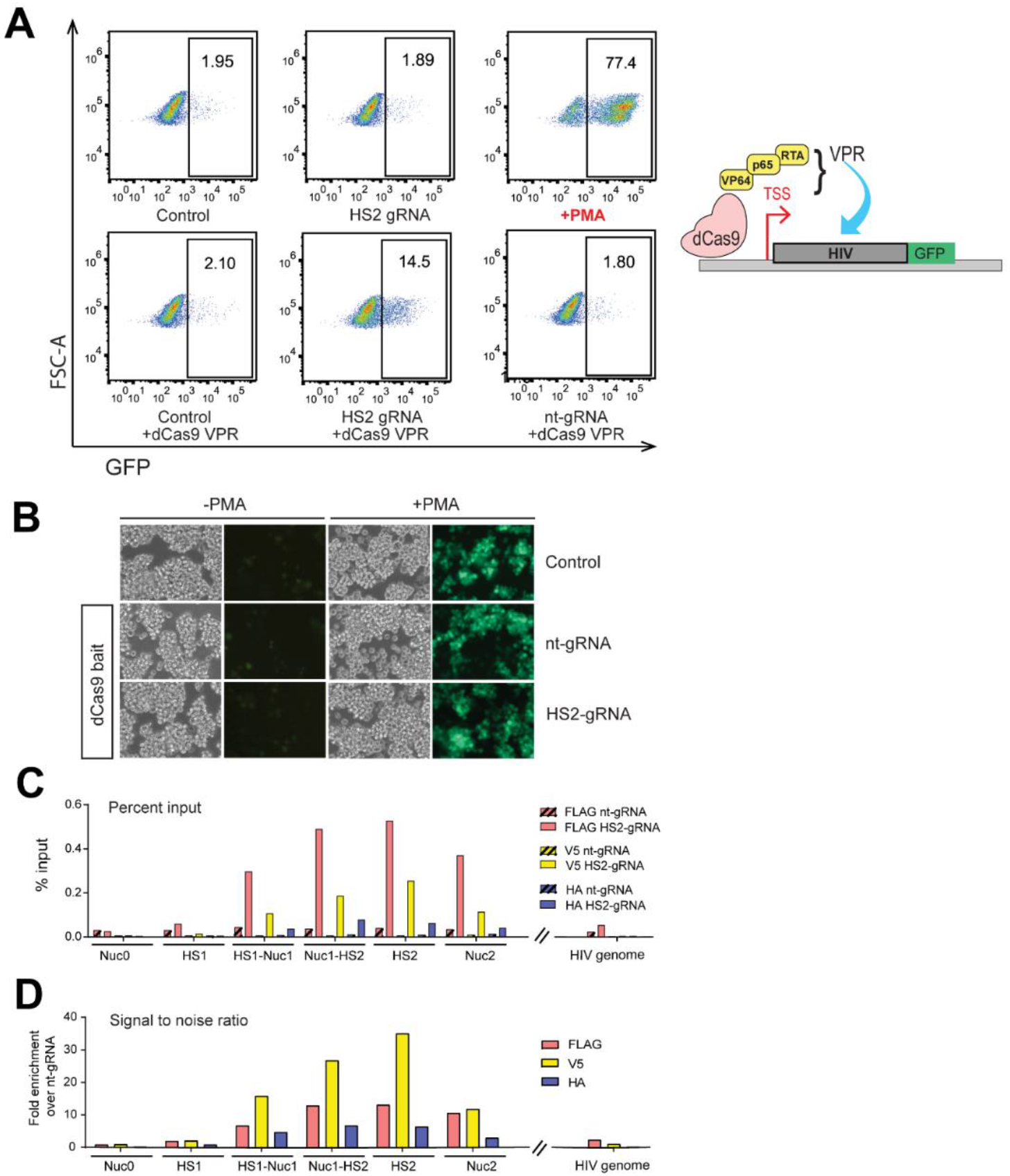
Characterization of the experimental system. (A) Functional validation of the gRNA designed against the HIV-1 5’LTR HS2 region (HS2-gRNA). The FACSs plots show J-Lat 11.1 cells examined by flow-cytometry at 72 hours after nucleofection with a dCas9 VPR construct to check for HIV-1 LTR dependent GFP expression, measured as % GFP positive cells. (B) Microscopy pictures (bright field and fluorescence) of control J-lat 11.1 cells, cells expressing dCas9 bait and a non-targeting gRNA, cells expressing the bait and a gRNA targeting the HS2 region. Cells have been examined in unstimulated (-PMA) and in the presence of 20nM PMA (+PMA) to assess reactivation capacity. (C) ChIP-qPCR experiments performed using different antibodies, conjugated to affinity beads, against the different synthetic tags (FLAG, V5, HA) of the dCas9 construct. Cells expressing the HS2 gRNA and control cells expressing a non-targeting gRNA (nt-gRNA) are compared. HIV-1 5’LTR sequences recovery is calculated as a percentage of the input. (D) From the experiment shown in (C), the signal to noise ratio of the experiment is calculated by dividing the ChIP-qPCR signal obtained in the HS2 gRNA expressing pool with the signal obtained in the nt-gRNA expressing pool. Data are represented as fold enrichment over the non-targeting gRNA signal.

**Figure S2.**
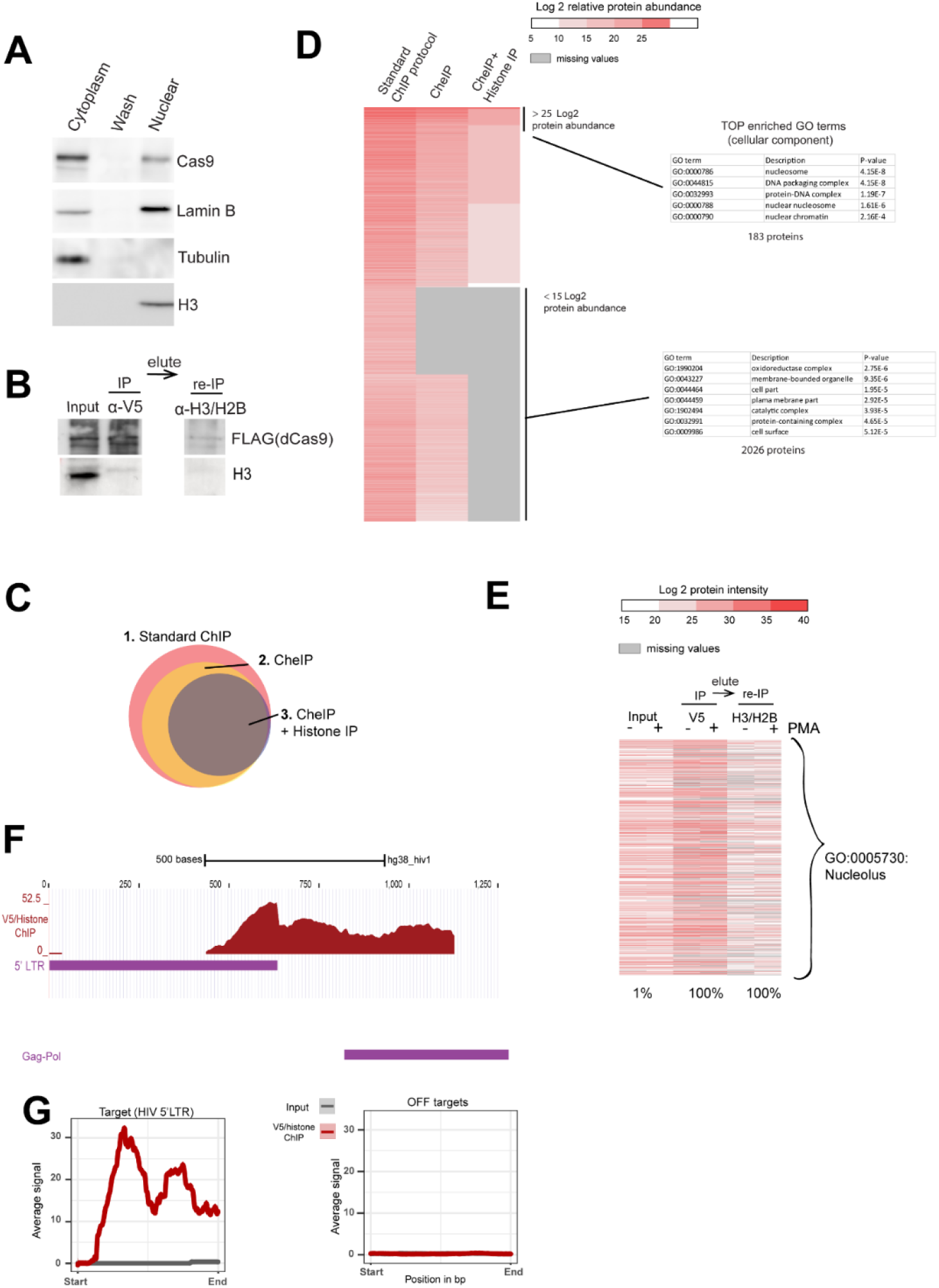
Characterization of the chromatin preparation protocol by mass spectometry and depletion of non localized bait. (A) Western blot analysis using antibody specific for dCas9 indicates localization of HA-V5-FLAG-dCas9 bait following a nuclear and nucleolar fractionation protocol. α-Tubulin is used as a cytoplasmic marker, Histone H3 is used as a chromatin marker while Lamin B is used as a nuclear marker. (B) Western blotting with anti Cas9 and anti-V5 antibody indicates relative presence of HA-V5-FLAG-dCas9 bait in the fractions used in the sequential ChIP experiments in Figure 1G. (C) Venn diagram showing the proportion of the number of hits identified in the different protocols (D) The heatmap shows a comparison between the protein content of a standard ChIP protocol, CheIP, and CheIP followed by a histone enrichment step (CheIP + histone IP) with H2B and H3 conjugated affinity beads. The colors represent the Log2 transformation of the proteins relative abundance. The protein relative abundance was calculated based on protein/peptide spectral intensity values and normalized to the total protein content (E) Heatmap displaying the content of nucleolar proteins (GO cellular compartment category GO:005730; nucleolus) at each step of the Catchet-MS purification pipeline used for isolation of the HIV-1 5’LTR. The colors range represents the represents the Log2 transformation of the proteins intensities scores. Values corresponding to the V5 based immunopurification are adjusted to the fraction of material analyzed by mass spectrometry, corresponding to 1:40 of the material used for the second, histone based (H2B/H3) immunopurification. Missing values are represented by grey lines. (F) ChIP-sequencing tracks of the V5/histone (H3/H2B) sequentially purified chromatin over the HIV-1 5’ LTR. (G) Average coverage profiles using ChIP sequencing reads mapped 500bp upstream and downstream of the peak center at the 5’LTR region of the HIV genome (’Targets’) to the respective coverage around the predicted off targets (’OffTargets’). ’Start’ denotes the starting base pair of the aforementioned 1kb region around the peak centers and ’End’ the ending base pair respectively.

**Figure S3.**
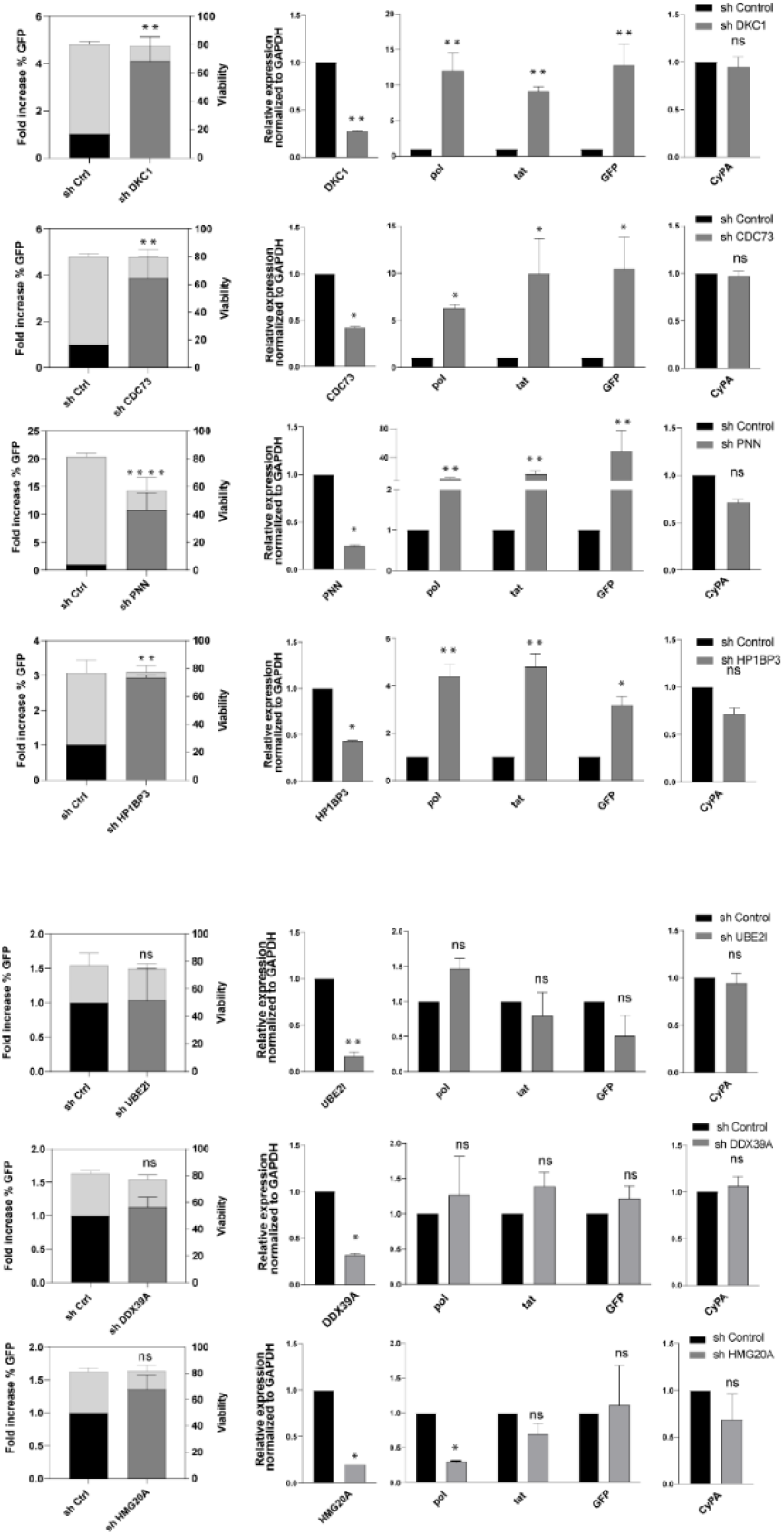
Functional validation of a selection of proteins bound downstream of the latent HIV-1 promoter. Functional validation of the hits associated with the repressed HIV-1 LTR. shRNA mediated depletion followed by Flow cytometry and RT-PCR. Statistical significance was calculated using ratio-paired t-test and multiple comparison t-test on Log2 transforment fold changes * – p < 0,05, ** – p < 0,01, *** – p < 0,001, ****– p < 0,0001

**Figure S4.**
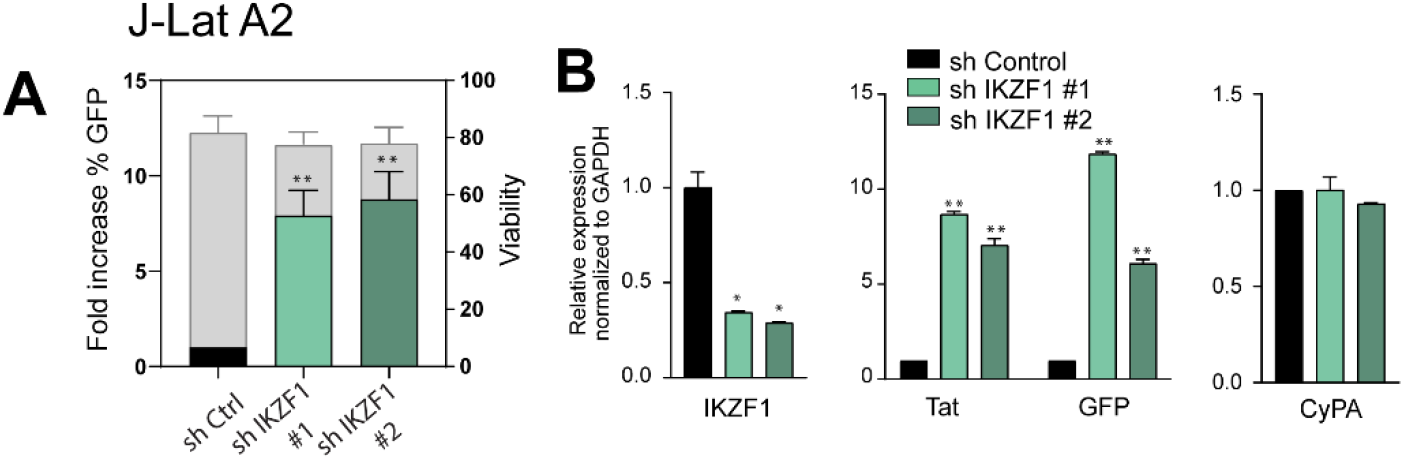
shRNA-mediated IKZF1 degradation in J-Lat A2 cells leads to latency reversal. (A) Bar plot showing the fold increase in % GFP positive cells (left y-axes) measured by FACS analysis, following IKZF1 depletion in J-Lat A2 cells with two different shRNA constructs (#1 and #2). The right y-axis represents the percentage of live cells. Data are the mean of two independent experiments (±SD). (B) qRT-PCR analysis measuring expression of HIV genes (pol, GFP, tat) in J-lat A2 cells transduced with scramble shRNA (sh Control) and sh IKZF1 #1 and #2. Data, normalized to GAPDH are represented as fold enrichment over sh Control and are the mean of three independent experiments (±SEM). Statistical significance was calculated using a ratio paired t test, * – p<0,05; **– p<0,01– ***p<0,001

**Figure S5.**
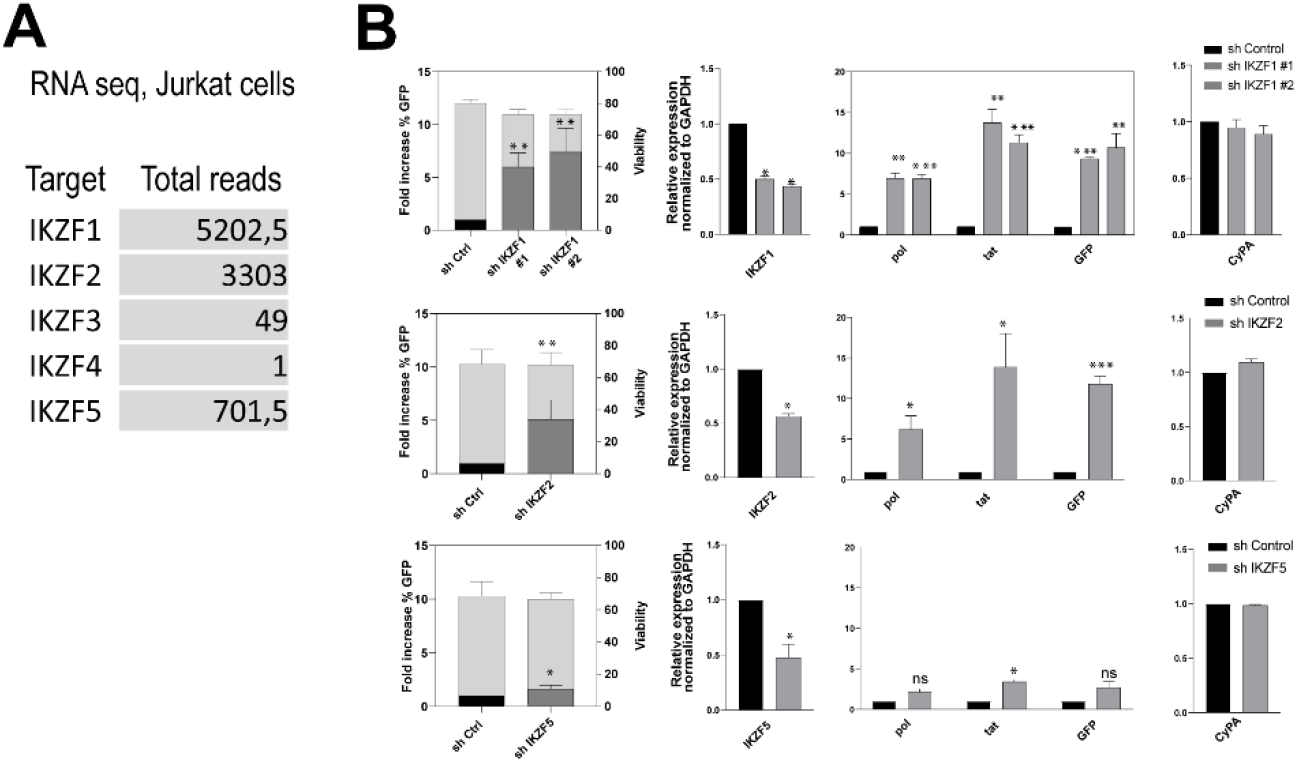
Expression of IKZF family members and their role in HIV-1 latency in J-Lat 11.1 cells. (A) Expression of IKZF family members in Jurkat cells. The panel shows the number of RNA seq reads in Jurkat cells. The data are published and available in Palstra et al., Science advances, 2018). (B) IKZF2 and IKZF5, the two prominent Jurkat cells expressed IKZF members were depleted from J-Lat 11.1 cells following shRNA-mediated transduction and GFP expression was examined by Flow cytometry and RT-PCR. Statistical significance was calculated using ratio-paired t-test and multiple comparison t-test * – p < 0,05, ** – p < 0,01, *** – p < 0,001, ****– p < 0,0001 .

**Figure S6.**
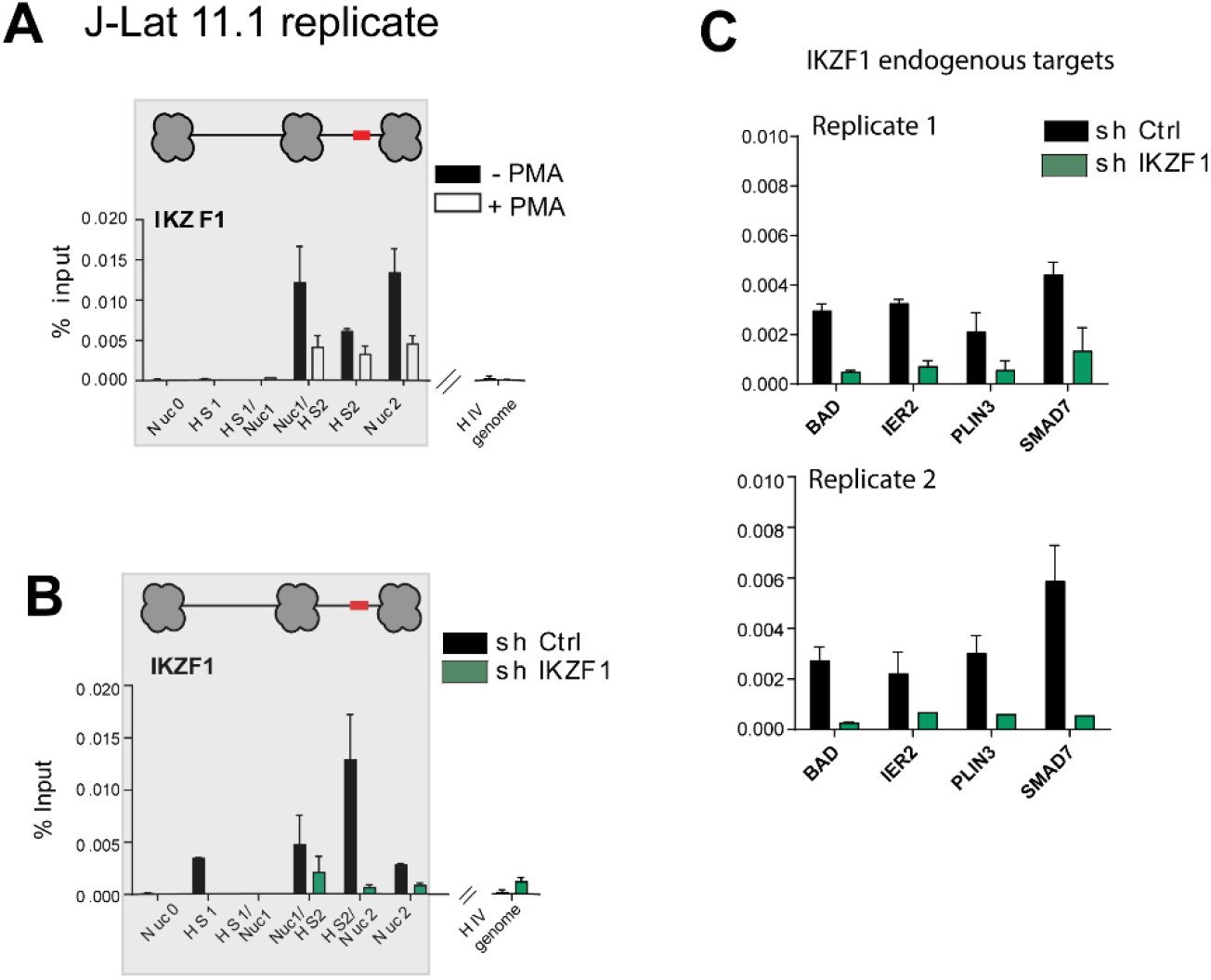
Carachterization of IKZF1 binding and chromatin state at the 5’LTR following treatment with PMA and shRNA-mediated IKZF1 degradation in J-Lat 11.1 cells. (A) Replicate ChIP-qPCR analysis with IKZF1 antibody in latent and PMA stimulated J-Lat 11.1 cells as indicated. Data are presented as % input, error bars represent the standard deviation (SD) of two separate real-time PCR measurements. (B) ChIP qPCR analysis using antibody against IKZF1 in J-Lat 11.1 cells transduced with scramble shRNA (shControl) and shIKZF1 probing binding to the HIV-1 5’LTR. Data is presented as % input, error bars represent the standard deviation (SD) of two separate real-time PCR measurements. (C) ChIP qPCR analysis using antibody against IKZF1 in J-Lat 11.1 cells transduced with scramble shRNA (shControl) and shIKZF1 probing binding of IKZF1 to endogenous IKZF target genes BAD, IER2, PLIN3 and SMAD7 (*46*).

**Figure S7.**
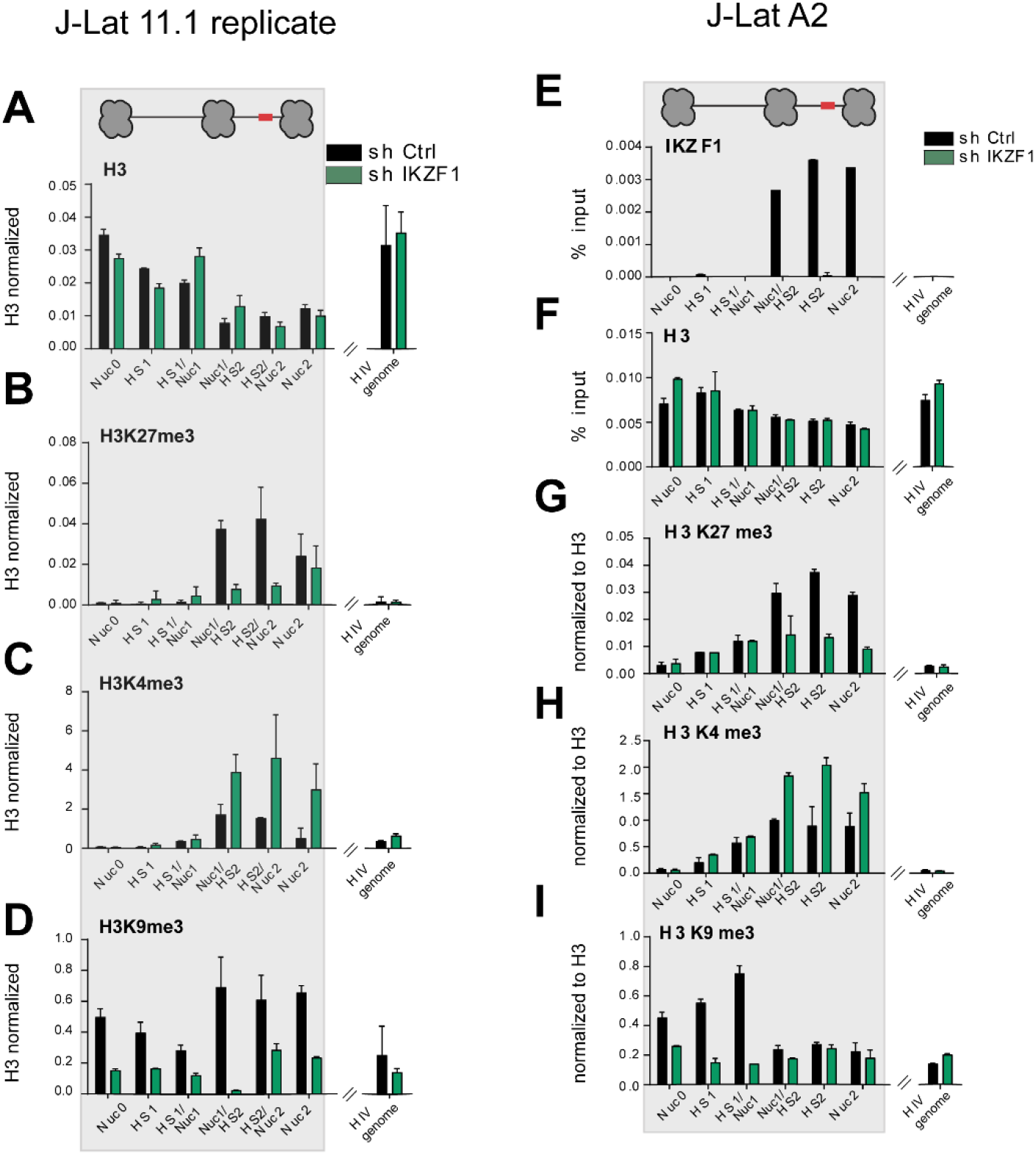
IKZF1, required for maintenance of HIV-1 latency in J-Lat 11.1 and A2 cells, binds downstream of the latent HIV-1 5’LTR and establishes of a repressive chromatin environment. (A-D) ChIP-qPCR using antibodies specific for distinct histone marks in J-Lat 11.1 cells transduced with scramble shRNA (shControl) and shIKZF1; total histone H3 (A), H3K27me3 (B), H3K4me3 (C), H3K9me3 (D). Total histone H3 data (A) are represented as % input mean (±SD), histone marks data (B-D) are expressed as fold change over H3 signal. Error bars represent the standard deviation (SD) of two separate real-time PCR measurements. (H) Representative agarose gels demonstrating range of DNA size resulting from sonication of chromatin used in the ChIP experiments presented in the manuscript. (E) ChIP qPCR analysis with IKZF1 antibody in J-Lat A2 cells transduced with scramble shRNA (shControl) and shIKZF1. Data are presented as % input, error bars represent the standard deviation (SD) of two separate real-time PCR measurements. (F-I) Histone marks ChIP qPCR analysis of the HIV-1 5’ LTR in J-Lat A2 cells transduced with scramble shRNA (shControl) and shIKZF1; (F) Total histone H3 (G) H3K27me3 (H) H3K4me3 (I) H3K9me3. Total histone H3 data (F) are presented as % input mean (±SD), histone marks data (G-I) are expressed as fold change over H3 signal. Error bars represent the standard deviation (SD) of two separate real-time PCR measurements.

**Figure S8.**
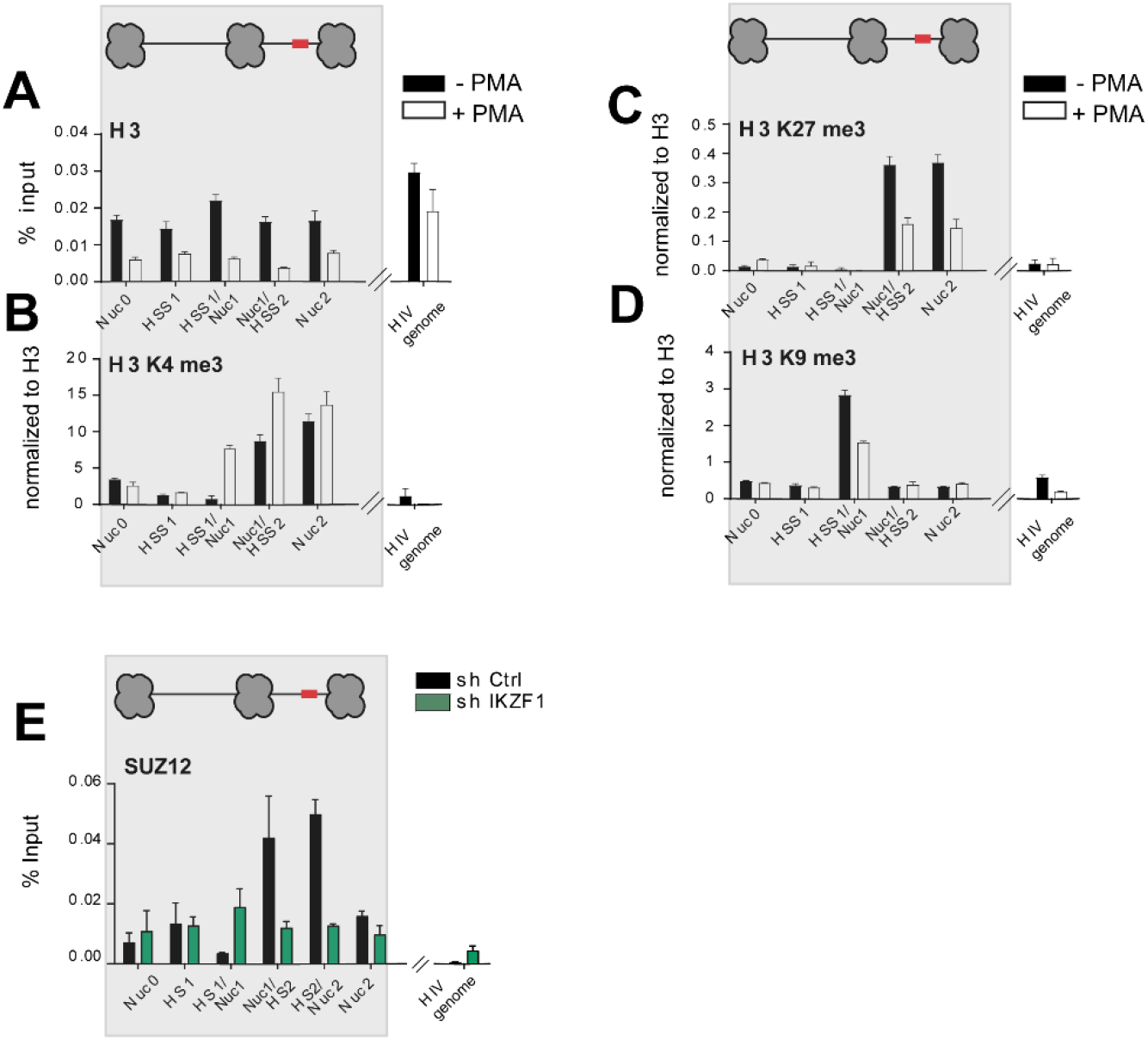
PMA stimulation leads to a loss of repressive chromatin marks and IKZF1 promotes PRC2 recruitment to the latent HIV-1 5’LTR. (A-D) ChIP-qPCR using antibodies specific for distinct histone marks in latent and PMA stimulated J-Lat 11.1 cells as indicated; total histone H3 (A), H3K4me3 (B), H3K27me3 (C), H3K9me3 (D). Total histone H3 data (A) are presented as % input mean (±SD), data corresponding to histone marks (B-D) are expressed as fold change over H3 signal. Error bars represent the standard deviation (SD) of two separate real-time PCR measurements. (E) ChIP-qPCR analysis with SUZ12 in J-Lat 11.1 cells transduced with scramble shRNA (shControl) and shIKZF1 at the HIV-1 5’LTR (Supplementary Figure S5B). Data is presented as % input, error bars represent the standard deviation (SD) of two separate real-time PCR measurements.

**Figure S9.**
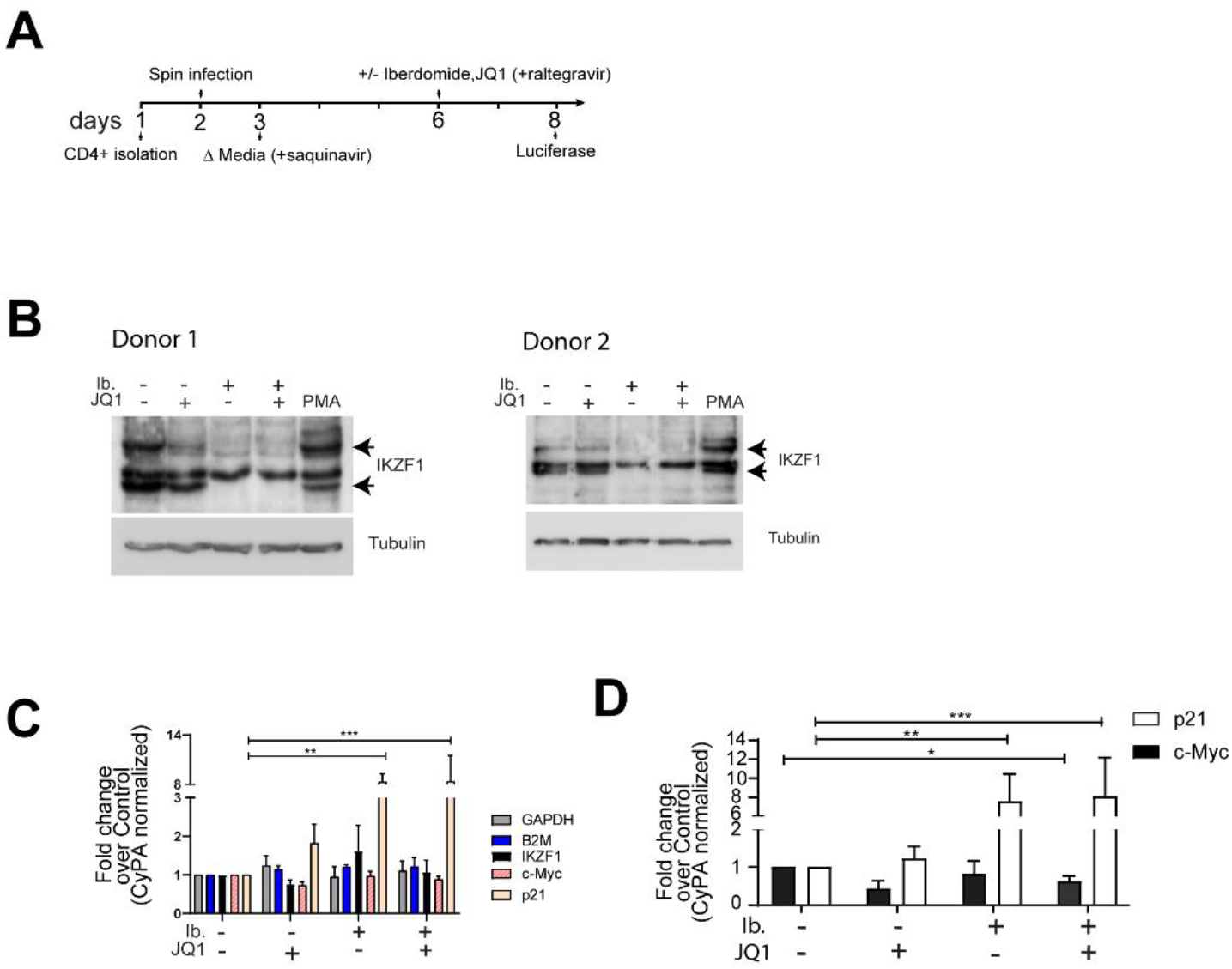
Iberdomide treatment of primary CD4+ T cells causes depletion of IKZF at the protein level and does not cause cytotoxicity or induce T cell activation. (A) Schematic representation of the protocol for HIV-1 latency establishment in primary human CD4+ T cells. (B) Western blot analysis using an antibody specific for IKZF1 indicates depletion of IKZF1 at the protein level in CD4+ T cells, upon treatment with iberdomide, JQ1 or a combination of both compounds as indicated for 24h. PMA is used as a control. α-Tubulin is used as a loading control. (C) qRT-PCR analysis of IKZF1 and IKZF1 targets p21 and c-myc upon treatment with JQ1, iberdomide, and their combination. RT-PCR was performed in primary CD4+ T cells isolated from three healthy donors. Data are represented as fold change (±SEM) over untreated and are normalized to Cyclophilin A (CyPA). B2M and GAPDH are used as housekeeping genes. Statistical significance was calculated using ratio-paired t-test * – p < 0,05, ** – p < 0,01, *** – p < 0,001. (D) qRT-PCR analysis of IKZF1 targets p21 and c-myc upon treatment with JQ1, iberdomide, or both compounds. RT-PCR was performed in primary CD4+ T cells isolated from 5 aviremic HIV-1 infected study partecipants. Data are represented as fold change mean (±SD) over untreated and are normalized with Cyclophilin A (CyPA). Statistical significance was calculated using ratio-paired t test * – p < 0,05, ** – p < 0,01, *** – p < 0,001.

**Figure S10.**
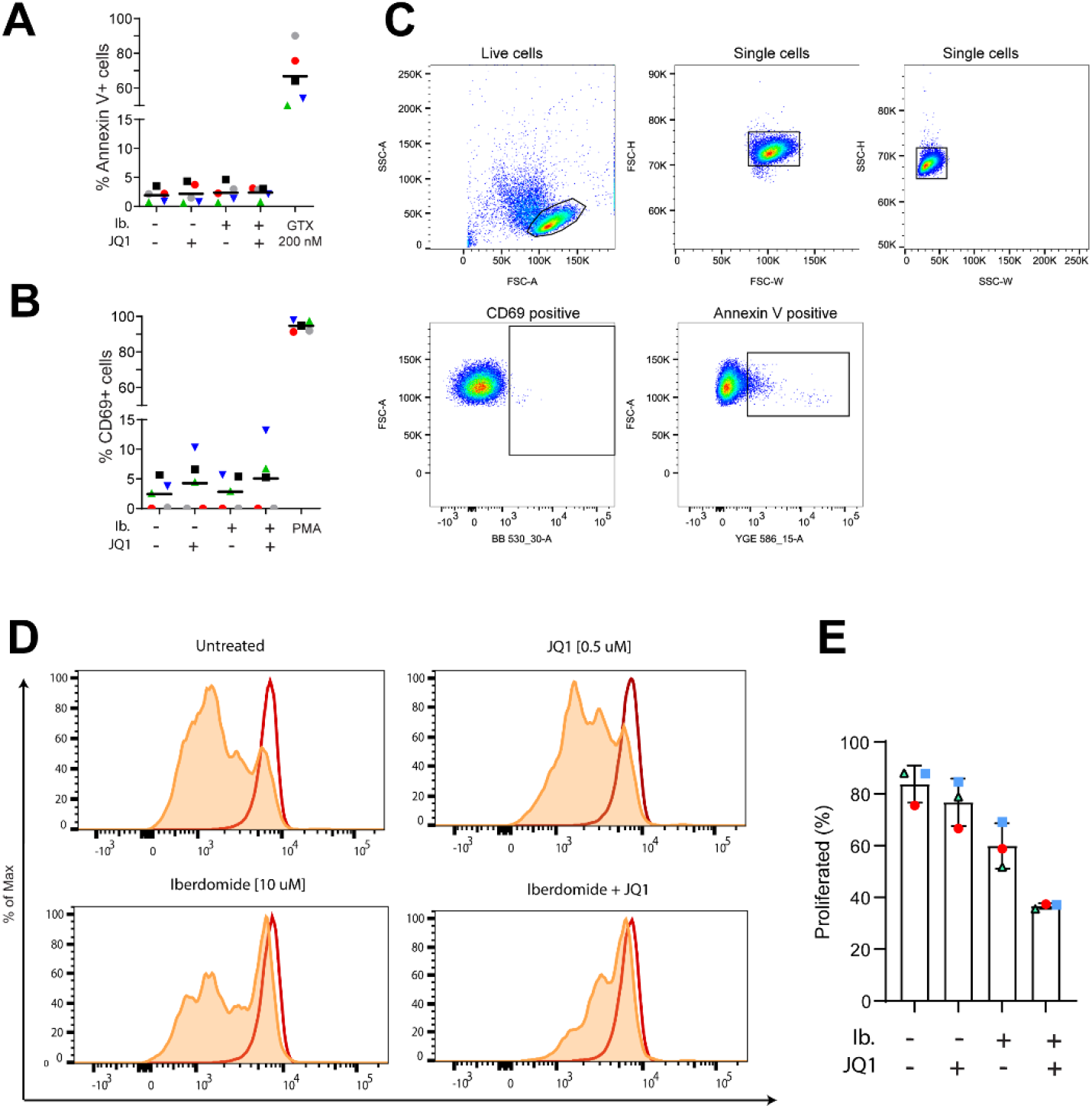
Iberdomide treatment causes reduction in proliferation capacity of CD8+ T cells but is not toxic to CD4+ T cells. (A) Percentage of cells expressing the Annexin V marker of apoptosis in primary CD4+ T cells treated with iberdomide for 48 hours, JQ1 and the combination of both compounds. Treatment with a toxic concentration of Gliotoxin (GTX) 200nM was used as a positive control. (B) Percentage of cells expressing the CD69 marker of cell activation in primary CD4+ T cells treated with iberdomide for 48 hours, JQ1 and the combination of both compounds. Experiments were performed in uninfected cells obtained from 5 healthy donors. Treatment with PMA was used as a positive control. Bars represent the average of experiments performed on samples deriving from two healthy donors. (C) Representative flow cytometry plots and gating strategy for annexin V and CD69 staining. (D) Representantive histogram of proliferative capacity of unstimulated or aCD3/CD28 stimulated CD8+ T cells in the presence or absence of LRAs. Cells were stained with a proliferation dye and analyzed 72 hours later by flow cytometry. Dividing cells show decreased intensity of proliferaiton dye as it becomes diluted upon cell division. (E) Percentage of proliferated CD8+ T cells from 3 healthy donors as described in C.

**Figure S11.**
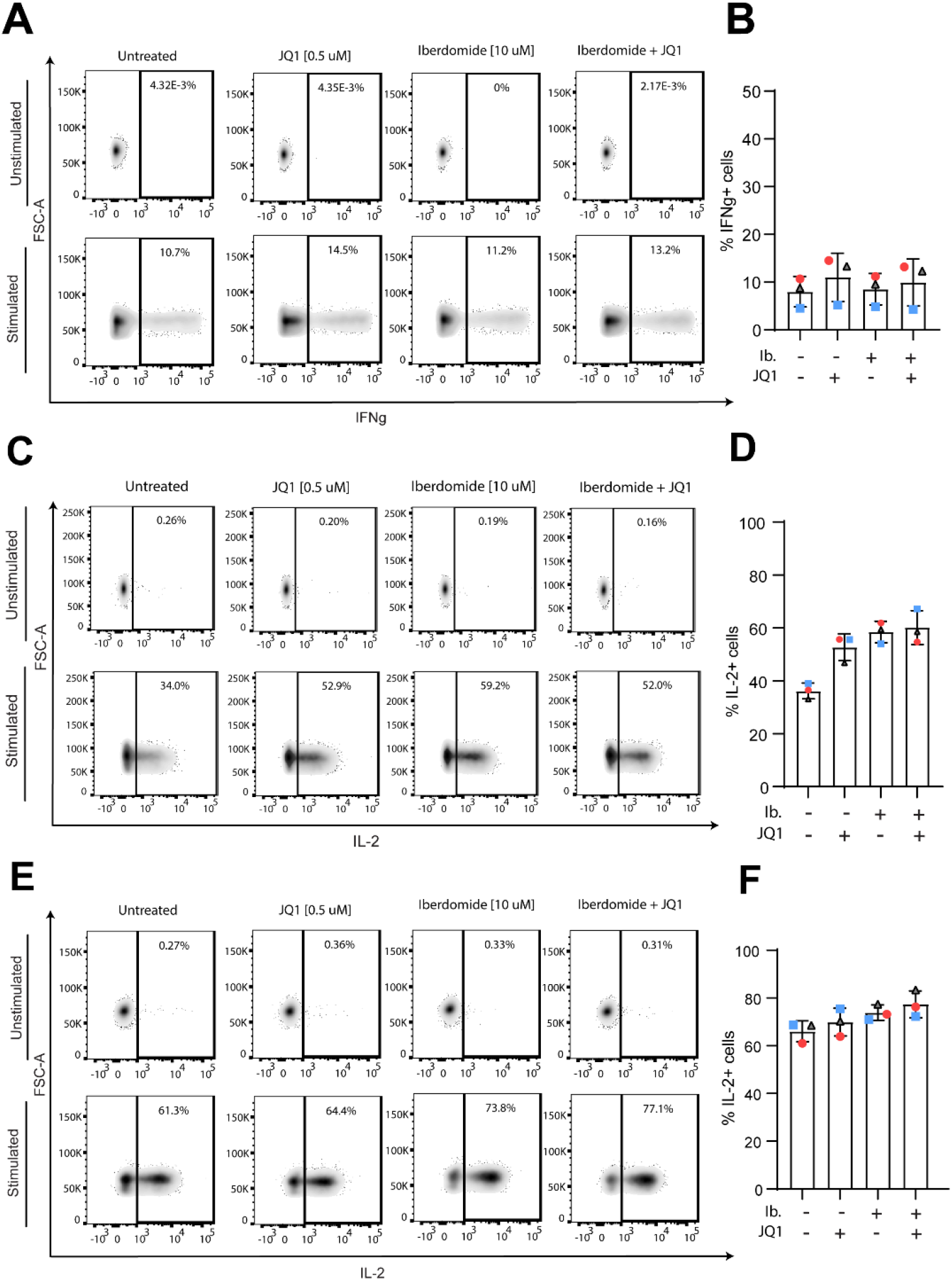
Effect of iberdomide treatment alone and in combination with JQ1 on T cell proliferation capacity and effector function. (A) Representative flow cytometry plots (left panel) of INF-g production analysis in unstimulated and stimulated primary CD4+ T cells. Cells were treated as indicated for 18 hours followed by PMA/Ionomycin stimulation for 7 hours in the presence of a protein transport inhibitor or remained unstimulated. IFNg production was assessed by intracellular staining and analyzed by flow cytometry. (B) Percentage of INF-g producing CD8+ T cells from 3 healthy donors as described in B. (C) Representative flow cytometry plots (left panel) of IL2 production analysis in unstimulated and stimulated primary CD4+ T cells. Cells were treated as indicated for 18 hours followed by PMA/Ionomycin stimulation for 7 hours in the presence of a protein transport inhibitor or remained unstimulated. IL-2 production was assessed by intracellular staining and analyzed by flow cytometry. (D) Percentage of IL-2 producing CD8+ T cells from 3 healthy donors as described in C. (E) Representative flow cytometry plots (left panel) of IL2 production analysis in unstimulated and stimulated primary CD8+ T cells. . Cells were treated as indicated for 18 hours followed by PMA/Ionomycin stimulation for 7 hours in the presence of a protein transport inhibitor or remained unstimulated. IL-2 production was assessed by intracellular staining and analyzed by flow cytometry. (F) Percentage of IL-2 producing CD8+ T cells from 3 healthy donors as described in E.

**Figure S12.**
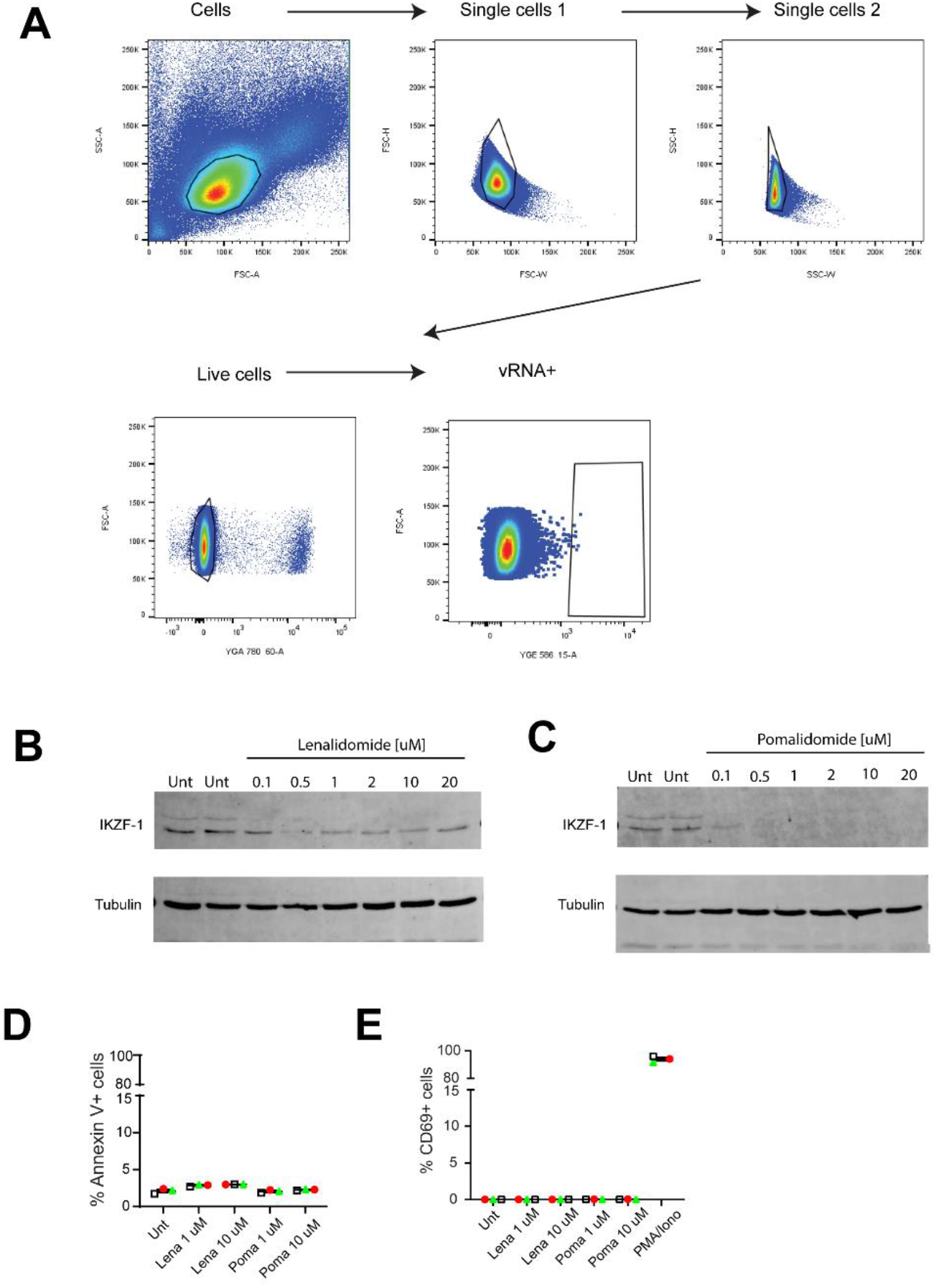
FISH-Flow analysis of pomalidomide and lenalidomide treated cells from HIV+ infected individuals and toxicity profile in CD4+ T cells from healthy donors. (A) Representative FISH-Flow flow cytometry plots and gating strategy for vRNA+ positive cells. (B) Western blotting shows protein levels of IKZF1 after 24 hours following treatment with Lenalidomide and Pomalidomide as indicated in primary CD4+ T cells. α-Tubulin is used as a loading control. (C) Percentage of cells expressing the Annexin V marker of apoptosis in primary CD4+ T cells treated with Pomalidomide and Lenalidomide for 24 hours. Experiments were performed in uninfected cells obtained from five healthy donors, represented by the dots. (D) Percentage of cells expressing the CD69 marker of cell activation in primary CD4+ T cells from 3 healthy donors as described in E. Treatment with PMA/Ionomycin was used as a positive control.

**Table S1.**
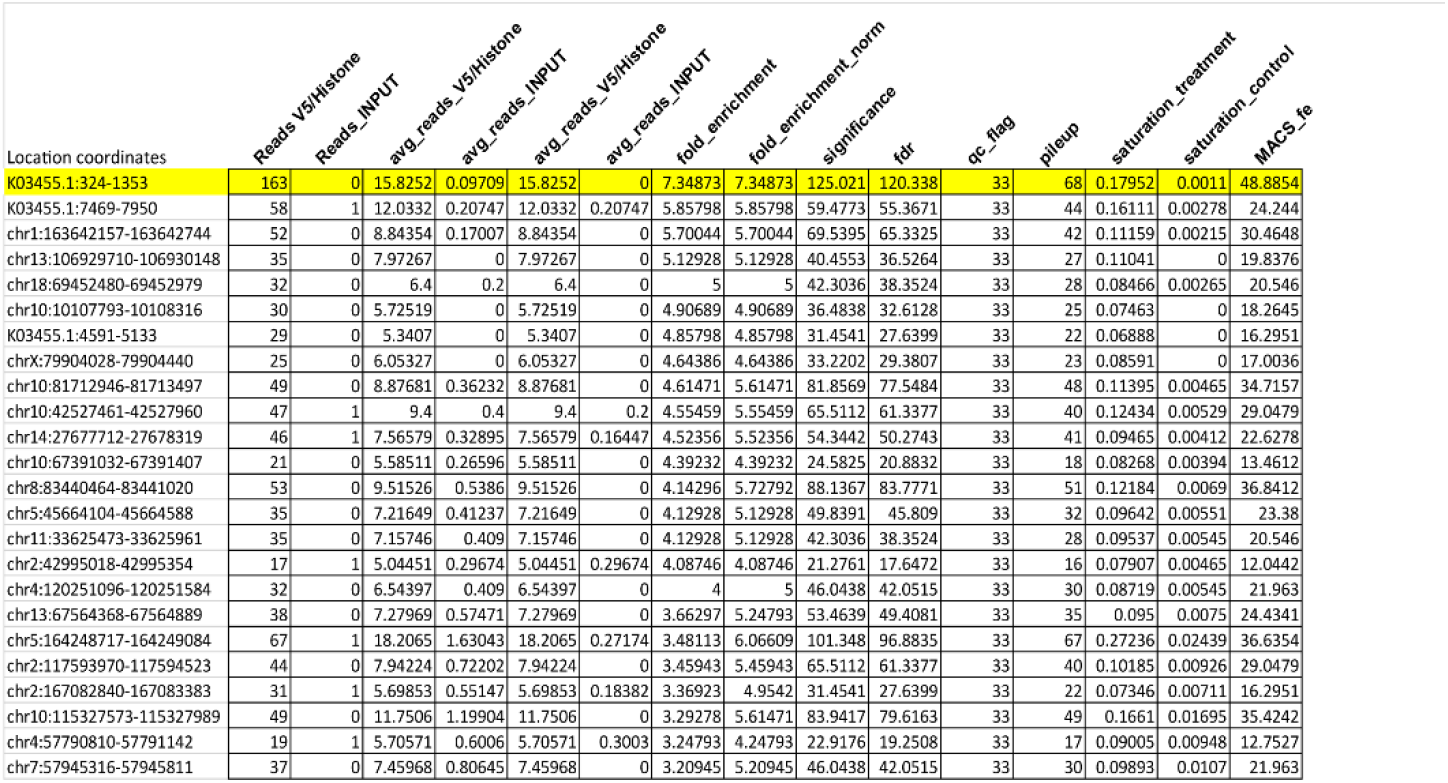
Supplementary Table 1 dCas9 V5/Histone ChIP sequencing peak calling summary.

**Table S2.**
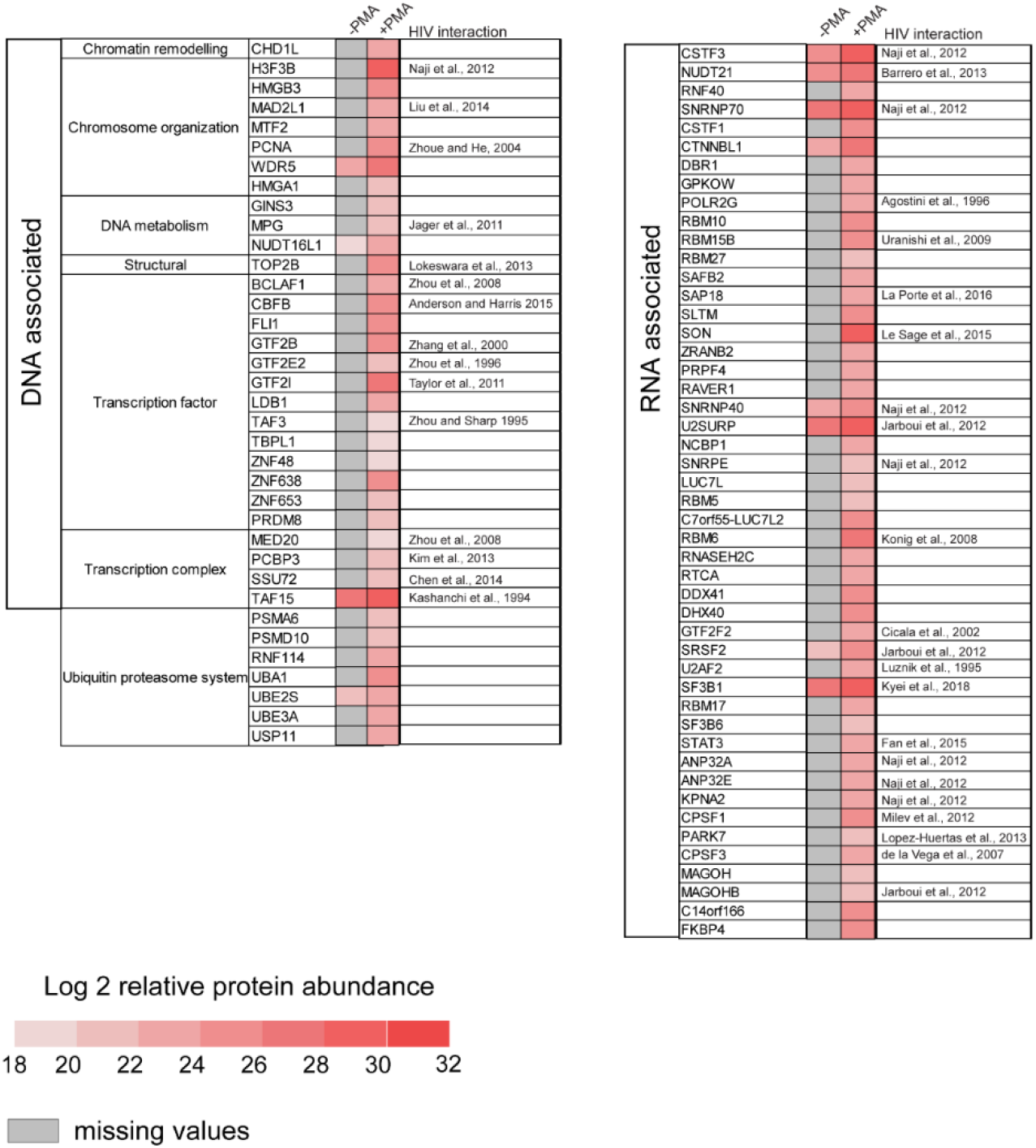
List of putative factors enriched on the active (+PMA) HIV-1 promoter. The table displays selected and functionally classified hits (n=84) identified by Catchet-MS in the +PMA state.

**Table S3.**
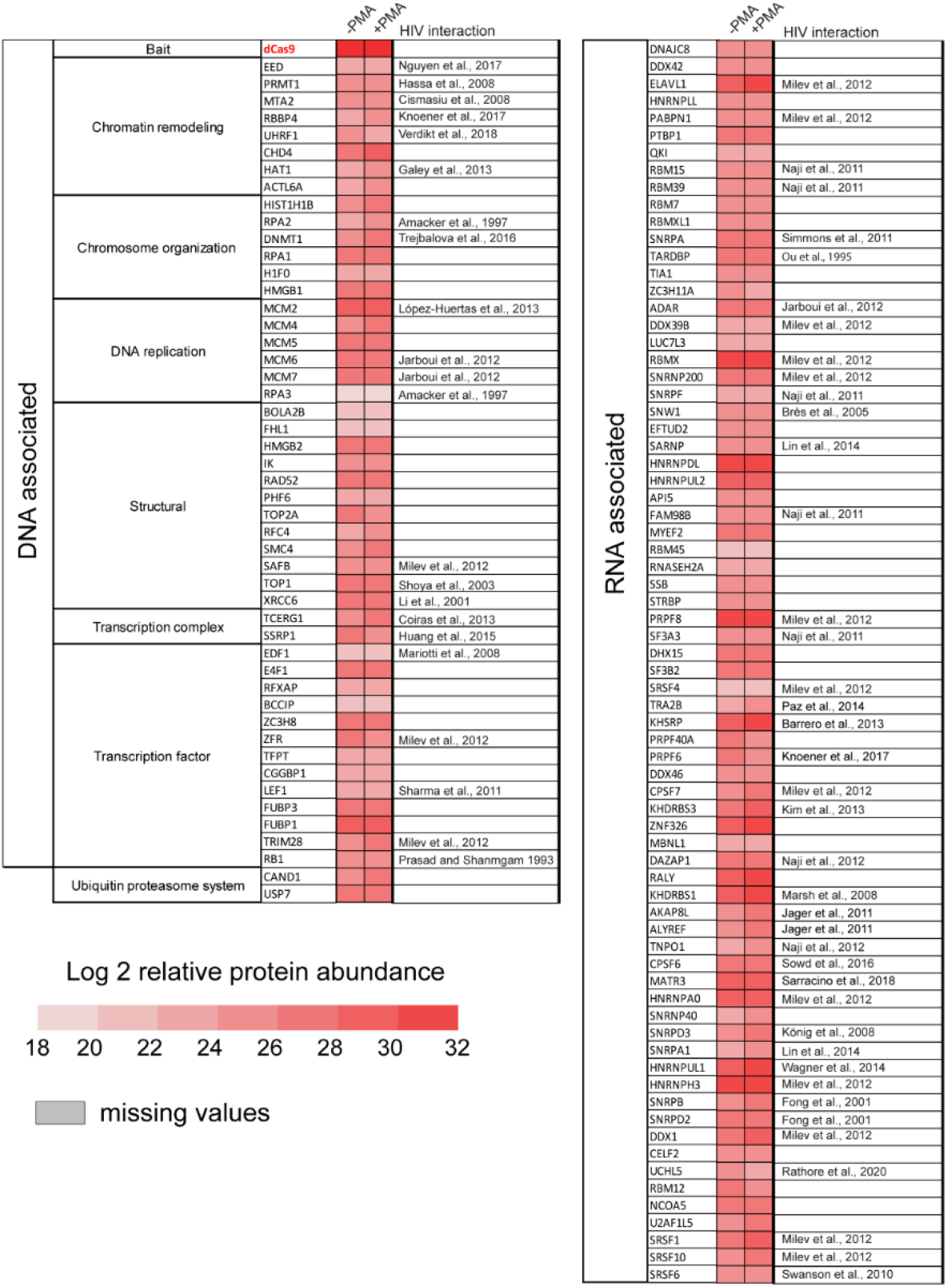
List of factors enriched on both the latent and active state of the HIV-1 promoter. The table displays selected and functionally classified hits (n=122) identified in both experimental conditions with similar scores, including the HA-V5-FLAG-dCas9 bait and potential non-specific binding contaminants.

